# Competition for MCM Loading at Origins Establishes Replication Timing Patterns

**DOI:** 10.1101/2020.09.20.305276

**Authors:** Livio Dukaj, Nicholas Rhind

## Abstract

Loading of the MCM replicative helicase onto origins of replication is a highly regulated process that precedes DNA replication in all eukaryotes. The number of MCM loaded on origins has been proposed to be a key determinant of when those origins initiate replication during S phase. Nevertheless, the genome-wide characteristics of MCM loading and their direct effect on replication timing remain unclear. In order to probe MCM loading dynamics and its effect on replication timing, we perturbed MCM levels in budding yeast cells and, for the first time, directly measured MCM levels and replication timing in the same experiment. Reduction of MCM levels through degradation of Mcm4, one of the six obligate components of the MCM complex, slowed progression through S phase and increased sensitivity to replication stress. Reduction of MCM levels also led to differential loading at origins during G1, revealing origins that are sensitive to reductions in MCM and others that are not. Sensitive origins loaded less MCM under normal conditions and correlated with a weak ability to recruit the origin recognition complex (ORC). Moreover, reduction of MCM loading at specific origins of replication led to a delay in their initiation during S phase. In contrast, overexpression of MCM had no effects on cell cycle progression, relative MCM levels at origins, or replication timing, suggesting that, under optimal growth conditions, cellular MCM levels not limiting for MCM loading. Our results support a model in which the loading activity of origins, controlled by their ability to recruit ORC and compete for MCM, determines the number of helicases loaded, which in turn affects replication timing.

## Introduction

The initiation of DNA replication is an exquisitely orchestrated and highly conserved process. Although the molecular biochemistry of initiation at individual origins continues to be elucidated in great detail (Bleichert, 2019), the mechanism governing the time at which different regions of the genome replicate has remained largely elusive (Boos and Ferreira, 2019). Nonetheless, replication timing has been implicated in a number of important processes, including gene expression, mutation rates, and chromatin structure, and has emerged as a reliable signature for many cancerous states (Gilbert et al., 2010; Goren and Cedar, 2003; Lang and Murray, 2011; Müller and Nieduszynski, 2017; Ryba et al., 2012). Which molecular mechanisms underlie the replication timing patterns, how they establish it, and what the consequences are when those mechanisms go awry is still a matter of intensive research (Rhind and Gilbert, 2013).

Origin licensing, the establishment of potential origins of replication in G1, is separated from replication initiation through a variety of biochemical mechanisms, which are particularly well-understood in budding yeast (Nguyen et al., 2001). In late M and G1, the Minichromosome Maintenance (MCM) complex, the heterohexameric catalytic core of the replicative helicase, composed of Mcm2-7, is shuttled into the nucleus with the help of Cdt1 (Tanaka and Diffley, 2002). In the nucleus, the Origin Recognition Complex (ORC), which loads MCM at origins, recognizes and binds origins at their ARS Consensus Sequence (ACS) (Eaton et al., 2010; Xu et al., 2006). After binding of its Cdc6 subunit, ORC recruits MCM-Cdt1 to the origins and loads MCM onto double stranded DNA (dsDNA), at which point Cdt1 is released and a second MCM is coordinately loaded (Ticau et al., 2015). The resulting MCM double hexamer (MCM-DH) establishes an origin that can be activated for replication through the action of the replication kinases CDK and DDK and accessory initiation factors (Labib, 2010).

Quantification of MCM components in budding yeast using a variety of techniques has yielded a wide range of values for the number of MCMs in each cell (Ghaemmaghami et al., 2003; Donovan et al., 1997; Lei et al., 1996). A meta-study using data from 21 publications reported median value estimates for MCM components that ranged from 2774-5360 molecules per cell for each of the components of the MCM helicase (Ho et al., 2018). This analysis suggests that MCM is present in cells at levels that allow for loading of many more than one MCM at each of the 400 or so origins that are present in the genome. Moreover, measurements of *in vivo* chromatin binding in budding yeast indicate that MCM is loaded at a much higher number than accounted by the loading of a single MCM-DH at known origins (Donovan et al., 1997).

Indeed, studies from various eukaryotic organisms have shown that MCM is loaded onto the genome at levels that are higher than necessary for replication under optimal conditions (Bowers et al., 2004; Edwards et al., 2002). However, the excess of loaded MCM appears to be necessary under conditions that lead to elevated levels of replication stress. In *Xenopus* egg extracts, excess MCM loading becomes essential for successful replication under conditions of replication stress, indicating that dormant, excess MCM molecules are activated to salvage stalled replication forks that result from replication stress (Woodward et al., 2006). In *C. elegans*, worms only become susceptible to low doses of the replication fork stalling drug hydroxyurea (HU) when their MCM levels are reduced (Woodward et al., 2006). Similarly, studies using human cells have suggested a mechanism where excess MCM hexamers that are loaded and not used during unperturbed replication becomes vital for successful replication after treatment with replication-stress-inducing agents (Ge et al., 2007; Ibarra et al., 2008). Altogether, this data suggests that eukaryotes load more MCM onto DNA in G1 than is used during the S phase of the cell cycle.

Although excess loading of MCM is important for a successful S phase in response to replication stress, it has also been implicated in contributing to the replication timing program under optimal conditions. Genome-wide ChIP experiments in budding yeast have shown that MCM is loaded at origins in different amounts, and that the level of MCM loading correlates with the time in S phases at which an origin fires (Das et al., 2015; Foss et al., 2020). In addition, mutation of the B2 element in ARS1, which that has been shown to be important for MCM loading at that origin, causes reduced MCM loading and a delay in replication timing (Zou and Stillman, 2000; Das et al., 2015). This data points to a model where the more MCMs that are available for activation at an origin in S phase, the higher the likelihood that the specific origin will initiate early. Indeed, this multiple-MCM model fits well with kinetic modeling of replication, which predicts that stochastic firing arising from differences in the availability of MCM in the presence of limiting initiation factors can give rise to the observed replication kinetics seen in budding yeast cells (Yang et al., 2010; de Moura et al., 2010).

Although cellular MCM pools seem to be in large excess to origins of replication in budding yeast, many origins still show low levels of MCM loading by ChIP-seq (Das et al., 2015). Moreover, although the molecular details of how MCM is loaded onto origins have been elucidated, it remains unclear how loading dynamics are regulated at individual origins genome-wide. In order to investigate the mechanisms regulating the level of MCM loading and to see what implications changes in such levels may have for replication timing, we performed experiments under conditions where cellular MCM levels were either increased or decreased. We aimed to test to two general models of the regulation of MCM loading: the ORC Activity model, in which the relative activity of ORC at individual origins regulates MCM occupancy, and the Origin Capacity model, in which the intrinsic capacity of each origin for MCM loading regulates MCM occupancy. Importantly, we performed both MCM ChIP-seq and replication timing assays in the same experiments, such that the density of MCM loaded in late G1 could be directly correlated to replication timing at specific origins. Furthermore, we used MNase ChIP-seq, which produces much higher resolution data than standard ChIP-seq, allowing us to localize where MCMs are loaded within origins with close to base-pair resolution. Using genome-wide MNase ChIP-seq and replication timing assays, we found that lowering cellular MCM levels caused differential loading of MCM, with MCM loading at some origins being sensitive to intermediate reductions of cellular MCM levels and other origins being largely resistant. Importantly, reduction in MCM levels correlated with delays in replication timing. Conversely, increasing the MCM levels in budding yeast had no effects on MCM loading onto DNA or replication timing in S phase, indicating that MCM loading under normal conditions is saturated. These results suggest that loading of MCM at origins in G1 is a dynamic process and that relative levels of helicase at origins are dependent on the origins’ abilities to recruit MCM as well as the levels of MCM pools in the cells. In particular, as described in the Discussion, our results are consistent with a hybrid model combining regulation by both ORC activity and origin capacity.

## Results

### Auxin-induced reduction in cellular MCM levels causes a dose-dependent reduction in viability and slower progression through S phase

To test the effects of reduced MCM pools on helicase loading and replication timing we employed the auxin-inducible degron (AID) system optimized for budding yeast (Nishimura et al., 2009; Nishimura and Kanemaki, 2014). In order to reduce the cellular pool of MCM, we tagged the endogenous copy of Mcm4 with the AID degron cassette (IAA17) followed by GFP at its C terminus. The MCM hexamer requires all six components for nuclear localization (Labib et al., 2001). In addition, degradation of any single component of the hexamer causes destabilization of the rest (Labib et al., 2001). Therefore, degradation of Mcm4 is expected to reduce the total cellular pool of MCM. As previously reported, continual exposure to increasing amounts of auxin caused a dose-dependent lethality in cells harboring degron-tagged Mcm4 (**Figure S1a**) (Nishimura et al., 2009). However, cells maintain viability under our experimental conditions (**Figure S1b**).

We determined the effect of lowered MCM levels on genome-wide loading and replication timing in synchronized cells. We synchronized the cells and reduced MCM levels prior to G1 in order ensure that MCM was not removed from active replisomes or pre-loaded origins. We first synchronized cells in metaphase using nocodazole, followed by release into α-factor for a G1 arrest. Mcm4 was degraded through the addition of auxin, first for 30 minutes during nocodazole arrest, then during the majority of the α-factor arrest. Following a 30 minute period of equilibration in the absence of auxin at the G1/S boundary, samples were collected for MCM ChIP-seq analysis, and the rest of the culture was released into S phase (**Figure 1a**). Immunoblots monitoring the levels of Mcm4-IAA17-GFP revealed that Mcm4 is efficiently degraded after addition of auxin in a dose-dependent manner (**Figure 1b**). Quantification of Mcm4 levels revealed that, compared to untreated cells, 30 µM auxin reduced Mcm4 levels to 14% of the endogenous levels, whereas 500 µM auxin reduced those levels to 7%. The observation that cells released onto plates containing the replication stress drug hydroxyurea (HU) showed a dose-dependent increase in sensitivity to auxin suggests that lowered Mcm4 levels reduce MCM function (**Figure S1b**). We performed flow cytometry analysis on S-phase-synchronized cells treated with 0 µM, 30 µM, and 500 µM auxin and found that degradation of Mcm4 also causes a dose-dependent delay in progression through S phase (**Figure 1c**). Together, these data show that the AID system can be used to reduce Mcm4 levels in a dose-dependent manner and that reduction of Mcm4 causes reduced viability, slower progression through S phase, and lower resistance to replication stress.

**Figure 1:**
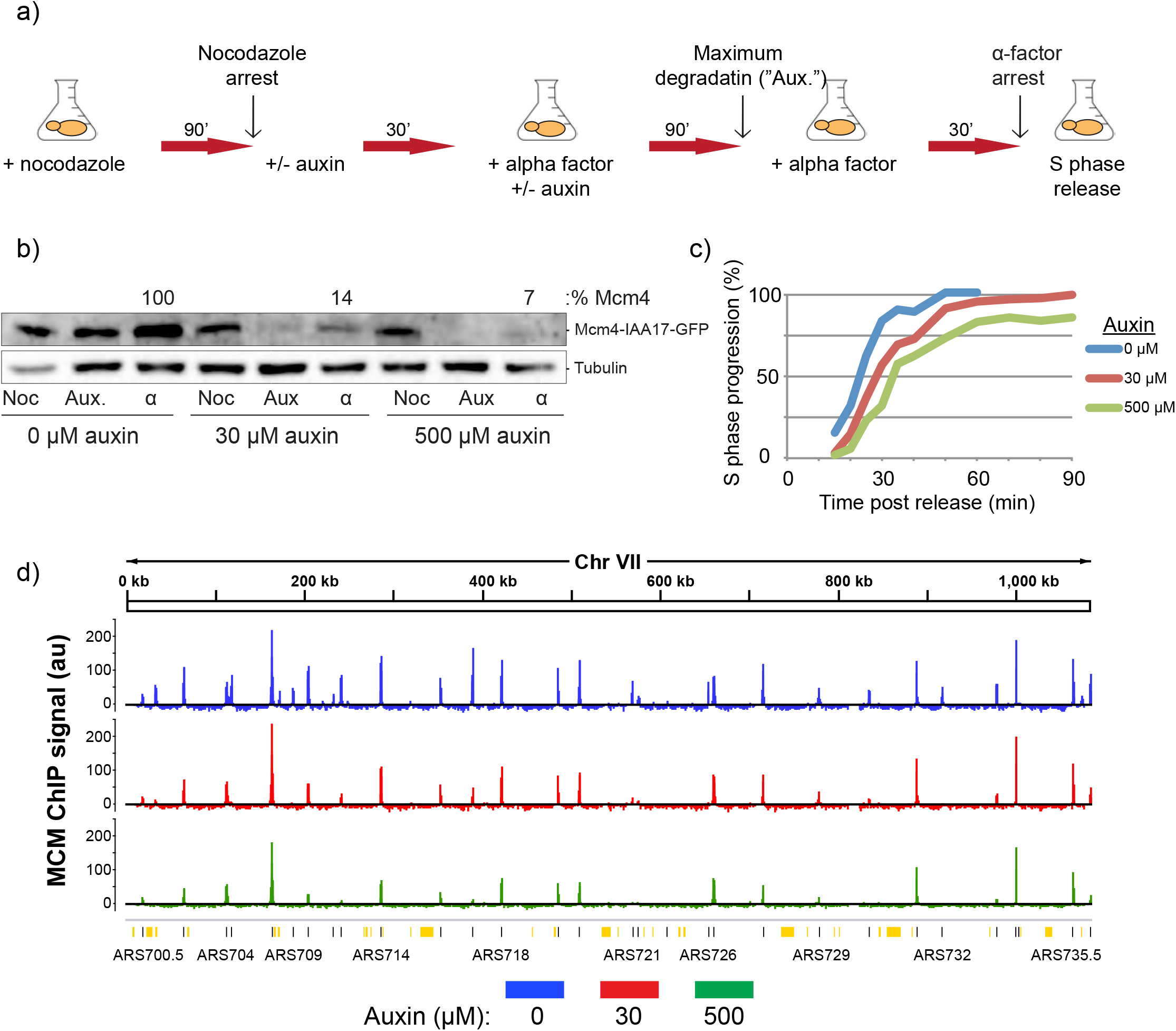
Reduction of MCM pools through auxin-mediated degradation of Mcm4. **a)** Experimental outline for MCM reduction experiments. yFS1059 cells were first synchronized with nocodazole, then released into α-factor for a G1/S arrest, prior to release into S phase. Flask symbol indicates filtering and release into new media, with additives as indicated. **b)** Western blot showing the levels of auxin-inducible cells (yFS1059) at the indicated time points of the experiment, probed using an anti-GFP antibody tracking GFP-tagged Mcm4. Noc = nocodazole arrest, Aux = maximum auxin degradation, α = α-factor arrest. Boxed value indicates quantitation of Mcm4-IAA17-GFP levels in α-factor arrested cells treated with 0 µM, 30 µM, or 500 µM auxin relative to levels of 0 µM auxin cells. **c)** Quantitation of flow cytometry data as cells progress from α-factor arrest through S phase. **d)** Normalized MNase-ChIP-seq coverage on Chromosome VII for non-induced cells. ARS-annotated origins are shown as black lines whereas potential, non-experimentally-confirmed origins are shown as orange lines/boxes (Siow et al., 2012).

### Reduced MCM levels cause a reduction in helicase loading at many, but not all origins of replication

Previous studies using ChIP have shown that MCM abundance at origins of replication varies throughout the genome (Xu et al., 2006; De Piccoli et al., 2012; Das et al., 2015). To test how lowered MCM pools affect the dynamics of helicase loading and abundance at origins of replication, we used α-factor-arrested cells to perform ChIP-seq using a polyclonal antibody against the full MCM hexamer on micrococcal nuclease (MNase) digested chromatin (Wal and Pugh, 2012). Mnase digestion has the advantage, relative to the standard ChIP approach of shearing by sonication, of cleaving the chromatin into much smaller fragments. Because the MCM double-hexamer complex protects about 70 bp of DNA, using MNase ChIP-seq, we can map MCM footprints with almost nucleotide resolution.

Fragments of the genome associated with MCM were pulled down, sequenced, and their abundance was normalized to read depth and no-immunoprecipitation input controls. Using this method, we found that MCM peaks localize to known origins of replication (**Figure 1d**), are reproducible (r = 0.97 - 0.99, **Figure S2b**) and correlate well with previously reported levels (r = 0.78 - 0.82, **Figure S2a**). Quantitation of origin peaks revealed changes to MCM loading throughout the genome in response to reduction of MCM levels (**Figure 2** and **Table S1**). Many origins exhibited a reduction in MCM abundance in auxin-treated cells, as seen by a shift away from the diagonal line in **Figure 2a**. The level of reduction was dose-dependent, as cells treated with 500 µM auxin showed greater reduction in MCM abundance than those treated with 30 µM auxin. Plotting the percent of MCM abundance that is lost in response to auxin treatment showed that some origins lose most of their MCM signal after 30 µM auxin treatment, while others are resistant to the changes in MCM levels even after 500 µM auxin treatment (**Figure 2b**). Origins that have lower levels of MCM in control cells are more prone to lose MCM signal in response to even intermediate levels of auxin treatment compared to origins that have high levels of MCM (**Figure 2b**). These results indicate that levels of MCM pools in cells affect loading at origins of replication throughout the genome in significant and dissimilar ways.

**Figure 2:**
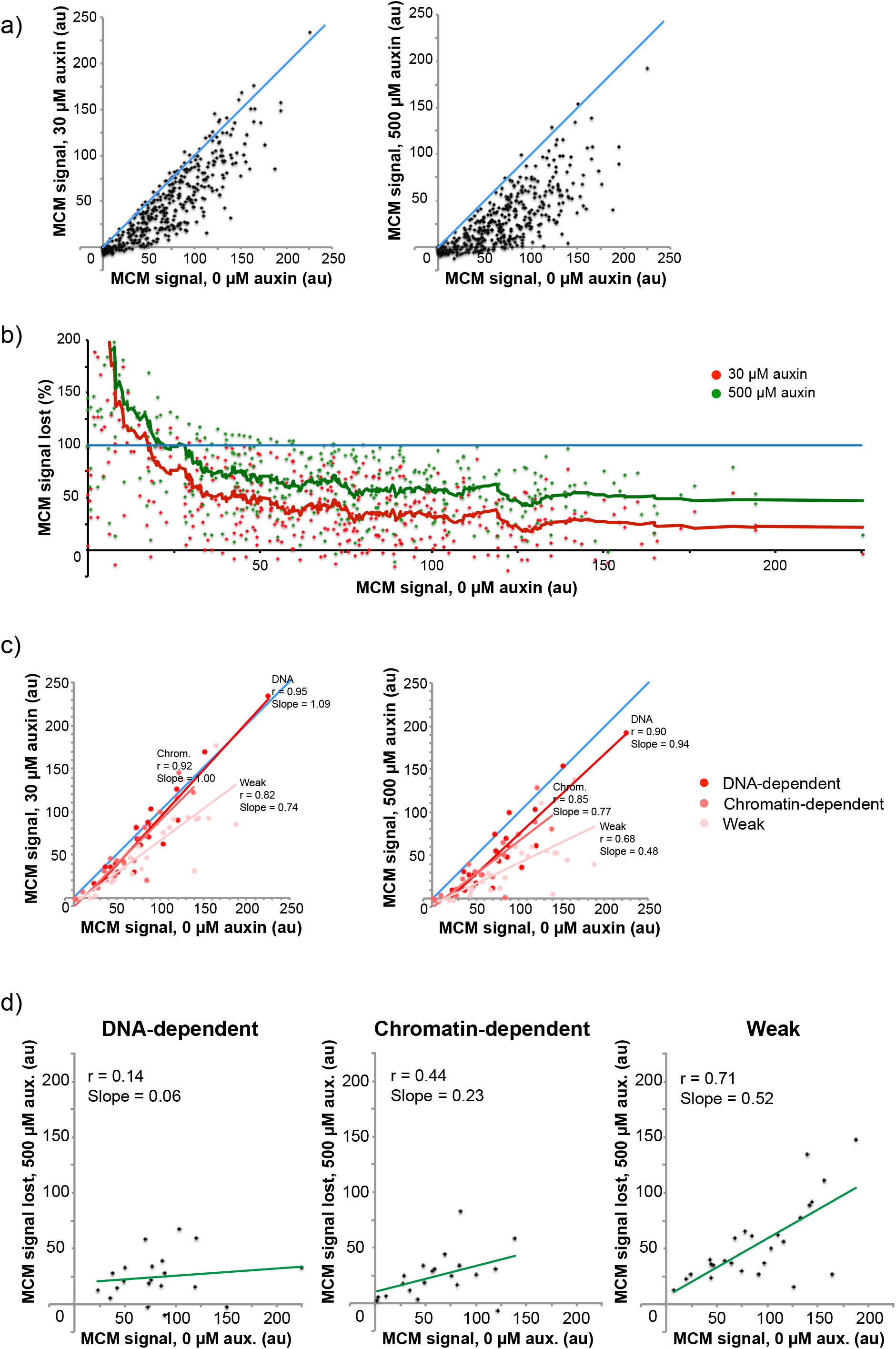
Reduction of cellular MCM levels causes a reduction in helicase loading at origins of replication. **a)** yFS1059 cultures were synchronized and released into S phase as shown in Figure 1a. MCM ChIP-seq signal at ARS-annotated origins from the indicated auxin treatments at the α-factor arrest point (x=y line drawn in blue for reference). **b)** Percent of MCM ChIP-seq signal lost upon treatment with 30 µM or 500 µM auxin. (20 pt moving average fit. y =100 line drawn in blue for reference). **c)** MCM ChIP-seq signal at 0 µM auxin treatment compared to 30 µM and 500 µM auxin for origins of replication characterized as ‘DNA-dependent’, ‘chromatin-dependent’, and ‘weak’ in Hoggard et al., 2013. **d)** MCM ChIP-seq signal for untreated cells plotted against the MCM ChIP-seq signal lost at the indicated auxin treatments at for the origins indicated in c).

To investigate the reason that reduced MCM levels delayed bulk S-phase progression, we examined its effect on the uniformity of the distribution of the majority of MCMs. We calculated the smallest number of origins required to account for 50% of the total MCM signal for each condition and measured the inter-origin distances between these major origins. If MCM pools are reduced and loading is decreased significantly for some origins, others would be expected to replicate larger tracts of the genome, leading to an increase in the time required to complete S phase (**Figure 1c**). As seen in **Table S2**, the smallest number of origins required to make up 50% of the MCM signal in each condition decreases as MCM pools are decreased (0 µM = 110, 30 µM = 85, 500 µM = 72), consistent with the observation that origins which load less MCM under control conditions lose a higher fraction of their MCM signal in response decreases in MCM pools. The *in-silico*-calculated average inter-origin distance between these top origins increases from 96 kb for 0 µM, to 118 kb for 30 µM, and to 136 kb for 500 µM auxin treatment, suggesting that they need to replicate larger tracts of the genome. In addition, a larger portion of origins replicate shorter tracts at 0 µM auxin, while a larger portion replicates longer tracts for 30 µM and 500 µM auxin treatments (**Figure S3**). Overall, this data indicates that, in response to reduced MCM pools, origins that load higher levels of MCM are responsible for replicating larger tracts of DNA, leading to an increase in the amount of time required to complete DNA replication.

### ‘Weak’ origins are more prone to reduced MCM loading

Previous analysis of various replication origins in budding yeast classified origins based on their affinity for ORC *in vivo*, as measured by ChIP-seq, and *in vitro*, as measured by gel shift assays (Hoggard et al., 2013). Origins for which high *in vitro* ORC affinity explained their high *in vivo* affinity were classified as DNA-dependent. Origins whose high *in vivo* affinity could not be explained by their *in vitro* affinity were classified as Chromatin-dependent. Origins that displayed low ORC affinity both *in vivo* and *in vitro* were classified as Weak. To gauge the effect of lowered MCM pools on loading dynamics between these origin classes, we compared MCM levels for cells treated with 0 µM auxin to those treated with 30 µM or 500 µM auxin (**Figure 2c**). The comparison shows that DNA-dependent and Chromatin-dependent origins maintain most of their endogenous levels of loading in response to auxin treatment and therefore are more resistant to changes in MCM pools compared to Weak origins. As expected, Weak origins lose more of their signal in response to reductions in MCM pools compared to DNA- and Chromatin-dependent origins (**Figure 2d**). This analysis suggests that origins with low ORC affinity are outcompeted by origins with higher ORC affinity (due to sequence or chromatin differences) when the pool of available MCM is reduced.

### MCM associates with flanking nucleosomes and the nucleosome-depleted region (NDR) of origins

Studies using MNase footprinting indicate that MCM associates with well-positioned nucleosomes flanking the ACS at origins of replication (Belsky et al., 2015). To assess the location of MCM signal in our data, we constructed strand-specific V plots centered at the ARS consensus sequence (ACS) (Eaton et al., 2010). V plots map the abundance of reads and their length relative to their genomic location. Mapping MCM ChIP-seq reads from control cultures shows that MCM is mostly found within 2-3 nucleosomes from the ACS, with signal decreasing with distance away form the ACS (**Figure 3a**). The signal includes MCM double hexamer-sized reads (∼68bp) as well as larger, nucleosome-sized reads (∼146bp) (**Figure 3a**). In contrast to previous genome-wide reports (Belsky et al., 2015), but in agreement with recent *in-vitro* cryo-EM structures (Miller et al., 2019), we also observe MCM signal in the nucleosome-depleted region (NDR) of origins. This observation was dependent on the degree of MNase digestion, as Replicate #1, which had a lower degree of digestion, lacked the smaller fragments, in agreement with similar studies involving transcription factors (**Figure S4**)(Henikoff et al., 2011).

**Figure 3:**
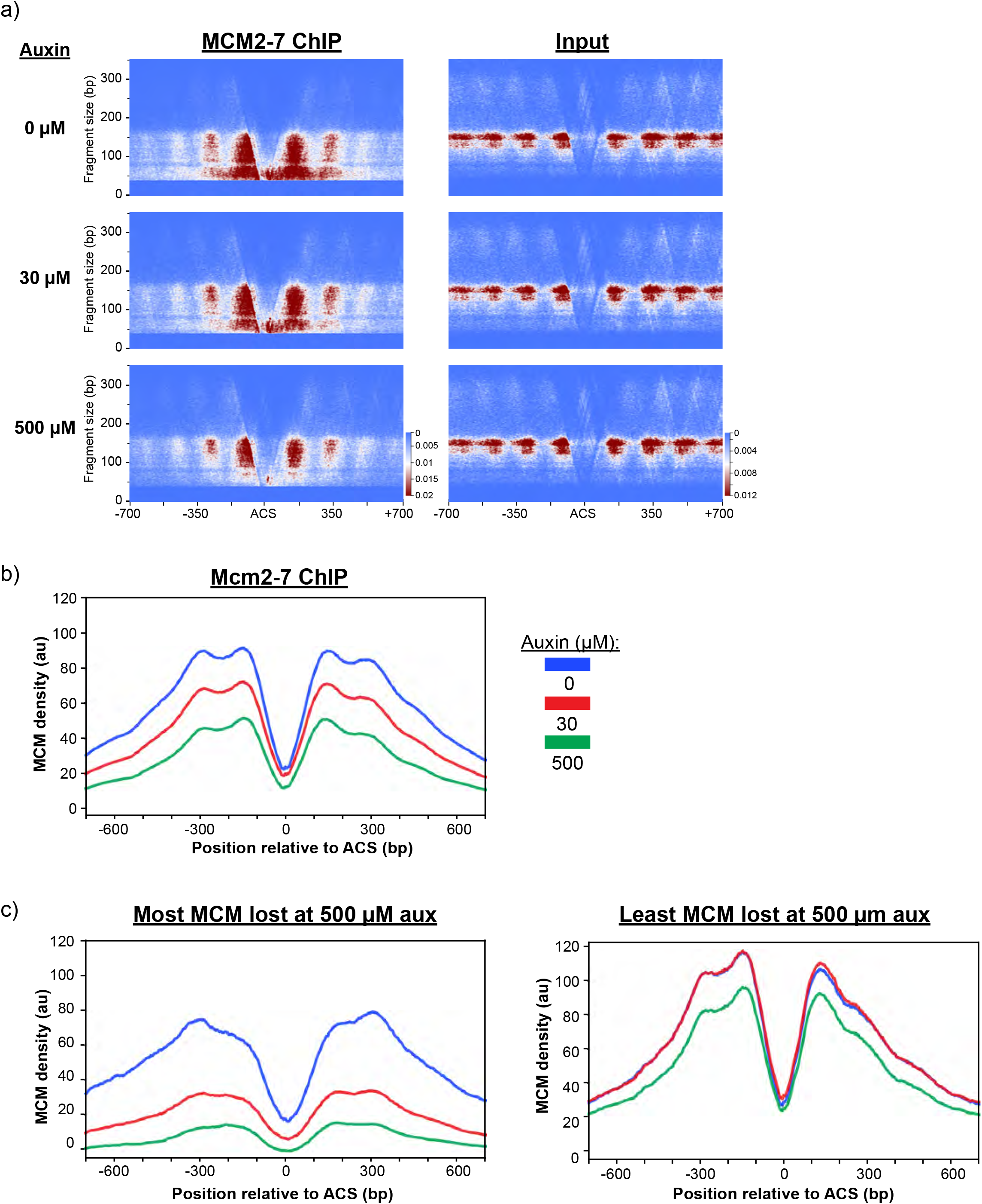
MCMs associate with the origin-adjacent nucleosomes and the NFR and show differential loading dynamics at origins. **a)** V plots of MCM MNase-ChIP-seq and input (non-IP) reads, plotted based on their length on a 700 bp window centered at 253 ACS-annotated origins (Eaton et al, 2010) (yFS1059 strain). **b)** Normalized MCM MNase-ChIP-seq density profiles centered at ACSs for the indicated auxin treatments. **c)** ACS-annotated origins were separated in quartiles based on the percent of signal lost in response to 500 µM auxin treatment compared to 0 µM auxin. The normalized coverage density profiles for the sets of origins displaying the largest reductions (‘Most MCM lost at 500 µM auxin’) and smallest reductions in MCM levels (‘Least MCM lost at 500 µM auxin’) are shown for all three treatments. Color scheme as in b).

In order to gauge changes in MCM signal around origins in response to reduction in MCM levels we constructed read density profiles. To do so, we used the midpoint of ACS-annotated origins and quantified read densities based on their genomic location around these origins (Eaton et al., 2010). As expected, there was a reduction in MCM ChIP-seq read density at origins in response to both 30 µM and 500 µM auxin (**Figure 3b**). In contrast, there was a much smaller but reproducible increase in read density for the +1 and −1 nucleosome positions for input DNA, indicating a possible increase in nucleosome occupancy as MCM loading is reduced at these origins (**Figure S5a**). However, in a 1kb window around the origins there was no significant difference in read density for input samples (r = 0.96-0.98, **Figure S5b**). Given the differential response of origins in response to auxin, we separated origins in quartiles based on the percent of MCM signal that they lost in response to 500 µM auxin. As seen in **Figure 3c**, origins which are least affected by MCM pool reduction tend to have higher levels of MCM loaded under control conditions and do not lose any signal in response to intermediate levels of auxin. On the other hand, origins which are most affected by reduced MCM pools tend to have less MCM loaded under control conditions and upon intermediate auxin treatment lose most of their signal. This data elucidates sets of origins that are distinct in how they are affected by MCM levels, identifying origins that are resistant to intermediate reductions in MCM levels and others that are sensitive.

ORC binding to the ACS and subsequent MCM loading is a directional process dependent on a ACS-site and a similar but inverted nearby sequence (Xu et al., 2006). To determine whether the directionality of MCM loading is maintained throughout G1 in relation to the dominant annotated ACS sites, we separated origins based on the magnitude of signal upstream or downstream of the ACS. The 253 origins tested were split nearly evenly between higher signal upstream or downstream of the annotated ACS (129 to 124, respectively; **Figure S6**). Notably, the MCM signal was largely independent of ACS directionality, as only 50% of higher-upstream-signal origins and 44% of higher-downstream-signal origins correlated with ACS directionality. This data supports the model where, once loaded, MCM is able to passively slide in either direction on double stranded DNA, consistent with previous *in vitro* findings (Remus et al., 2009; Miller et al., 2019).

### Auxin-induced reduction in MCM loading causes significant delays in replication timing at corresponding origins

Reduction of MCM pools via auxin-induced Mcm4 degradation caused significant changes to MCM loading at origins. To test whether the changes in MCM abundance have an effect on replication timing, we measured replication timing in cells released into S phase after treatment with 0 µM, 30 µM, or 500 µM auxin. Briefly, using the sync-seq approach, these experiments monitor replication timing by measuring genome-wide copy number at multiple points after synchronous release into S phase (Müller et al., 2014). From this data, the parameter T_rep_ (the time at which 50% of cells have replicated a specific locus) can be extracted. **Figure 4a** shows the T_rep_ profile of Chromosome V as well as the corresponding ChIP-seq profiles for treated and untreated cells. T_rep_ values for cells expressing control levels of MCM correlate well with values obtained from previously published studies (r = 0.91, **Figure S2a**) (Müller et al., 2014). Overlaying of T_rep_ profiles reveals that some origins that are active in untreated cells become inactive in auxin-treated cells. Loss of origin activity from these origins correlates with a decrease in MCM signal by ChIP, as is evident for ARS512 and ARS520 (**Figure 4a**, black boxes). Comparison of T_rep_ values between untreated and auxin-treated conditions indicates that reduction of MCM levels leads to delays in replication timing for a large number of origins and that these delays increase with higher auxin concentrations (**Figure 4b**). Together, this data demonstrates that reduction of cellular pools of MCM and subsequent reduction of MCM levels loaded at origins has direct implications for replication timing at specific origins in yeast.

**Figure 4:**
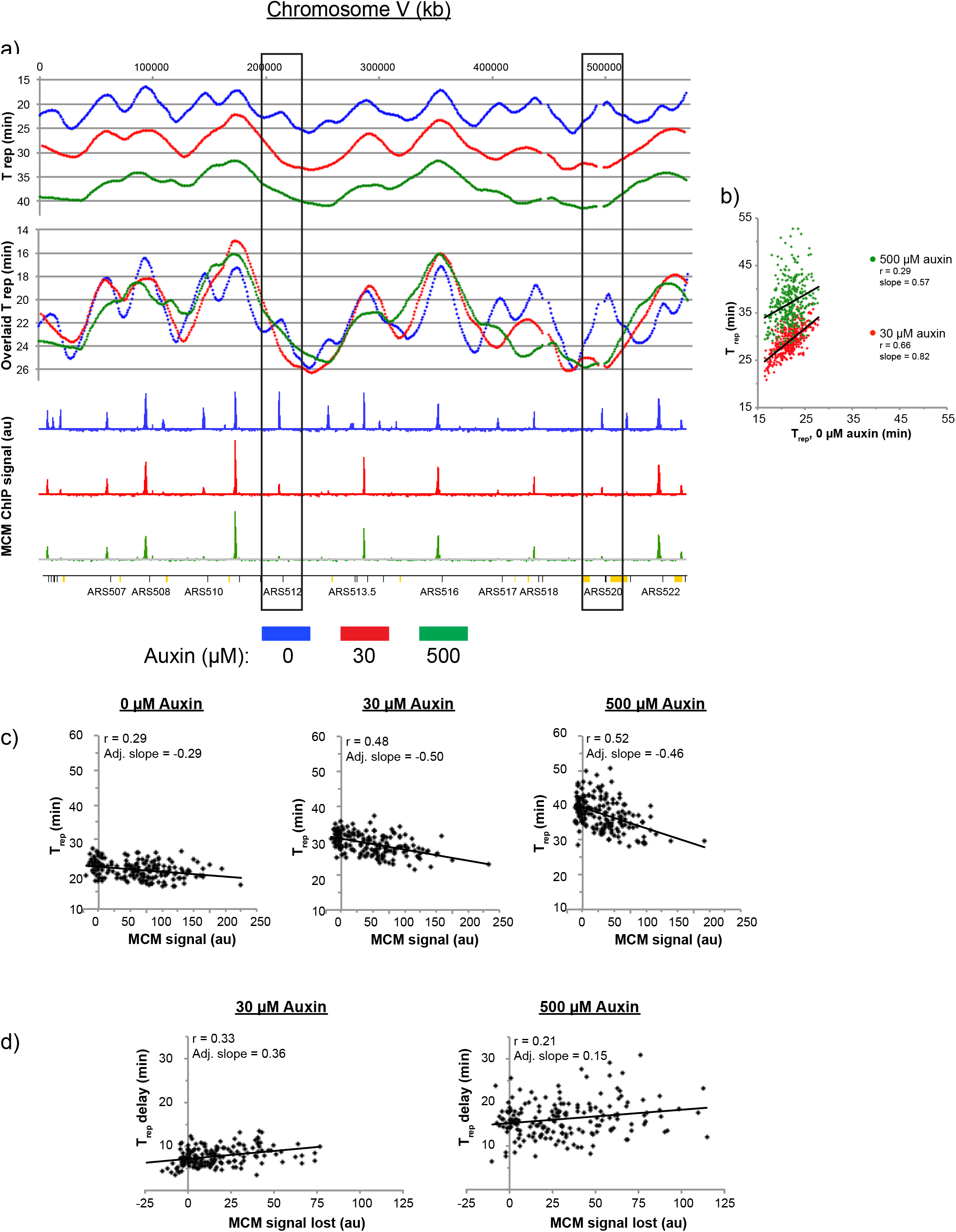
Delays in replication timing correlate with reductions in MCM levels. **a)** Chromosome V T_rep_ values for the 0 µM, 30 µM, and 500 µM auxin treatments (yFS1059 strain) was calculated in 1kb windows, LOESS-smoothed, and plotted based on their genomic location. “Overlaid T_rep_” values have been overlaid in order to compensate for delays in replication initiation and facilitate comparison between samples. Black boxes indicate origins that display strong correlation between the loss of MCM signal and delay in replication timing. ARS-annotated origins are indicated in black dashes on the bottom axis, whereas orange boxes denote unconfirmed origins from OriDB (Siow et al., 2012). **b)** T_rep_ values of ARS-annotated origins for untreated cells are plotted against values observed for 30 µM and 500 µM auxin treatments. **c)** Correlation between MCM signal and replication timing for origins that are not known to be affected by specific replication timing control factors (Das et al. 2016). **d)** Correlation between delay in replication timing as a result of auxin treatment and reduction in MCM signal, for origins plotted in c).

The replication timing of many origins throughout the budding yeast genome is controlled by specific mechanisms involving trans-acting factors and chromosomal location, such as Rpd3, Fkh1, and Ctf19 binding, or telomere proximity, (Knott et al., 2009; Lian et al., 2011; Knott et al., 2012; Natsume et al., 2013). We measured T_rep_ values and MCM abundance for the 43% of origins of replication that exclude those known to be affected by specific mechanisms of origin regulation (Das et al., 2015). Comparison of T_rep_ values and MCM abundance for these origins under endogenous conditions indicated a small negative correlation between MCM abundance and timing of replication, suggesting that origins with higher MCM levels replicate earlier in S phase (**Figure 4c**, 0 µM auxin). The correlation increased and the slope became more negative in conditions where MCM levels were reduced via auxin (**Figure 4c**, 30 µM and 500 µM auxin). Furthermore, comparing changes in T_rep_ with changes in MCM abundance as a result of auxin treatment revealed a positive correlation, indicating that larger decreases in MCM loading correlate with stronger delays in replication timing (**Figure 4d**). Taken together, these results suggest that decreases in MCM loading lead to delays in replication timing at origins of replication.

### Cells overexpressing MCM are viable

Reduction of MCM pools leads to significant changes in MCM loading and replication timing. In order to investigate the effects of the converse condition of increased MCM pools, we employed the galactose overexpression system. The six genes encoding MCM were expressed from bidirectional Gal-1,10 promoters integrated into the genome, in addition to their endogenous copies (Ticau et al., 2015). The overexpressed copy of Mcm7 was tagged at its C terminus with GFP in order to monitor its overexpression levels. To test whether cells overexpressing MCM are viable, we performed serial dilution assays on YP-Raffinose or YP-Galactose plates. Overexpression of the complete hexamer, with or without the added GFP tag on Mcm7, did not affect cell viability (**Figure S7a**). However, overexpression of the MCM hexamer in conjunction with the loading factor Cdt1, but not overexpression of Cdt1 alone, reduced viability. This data shows that Mcm2-7 overexpression by itself is not lethal and can be used to increase cellular pools of MCM helicase.

### Overexpression of MCM during a G1 arrest does not affect progression through S phase

We assayed whether overexpression of MCM leads to changes in helicase loading prior to S phase, and whether such changes affect replication timing. We synchronized cells in metaphase using nocodazole then released them into media containing α-factor for synchronization at the G1/S boundary (**Figure 5a**). In addition to α-factor, the media was supplemented with galactose or raffinose, to induce or repress Mcm2-7 overexpression, respectively. Induction of MCM was monitored via western blot using a polyclonal antibody against MCM. After two hours, a ∼5-fold overexpression of the hexamer was observed (**Figure 5b**). A portion of the α-factor arrested culture was collected for ChIP-seq analysis while the rest was released into S phase to monitor replication timing. As seen in **Figure 5c**, cells overexpressing MCM during a G1 arrest show a slight delay in replication initiation in S phase. However, a control strain that did not overexpress MCM showed similar results, indicating that the delay was likely due to the switch in carbon source (**Figure S7a**). Therefore, overexpression of MCM does not appear to cause any significant changes to cell cycle progression.

**Figure 5:**
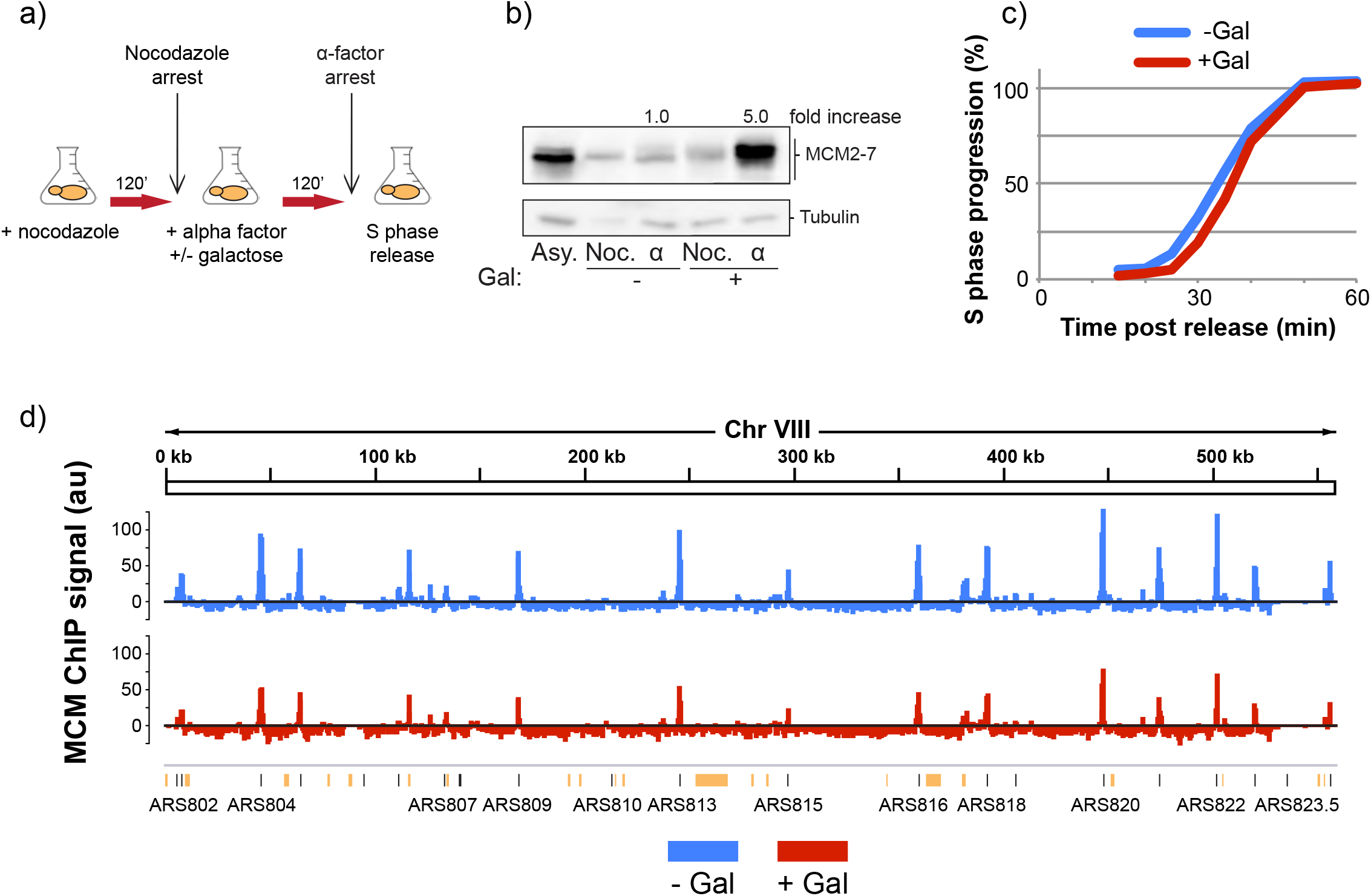
Galactose-induced MCM overexpression. **a)** Experimental outline for MCM overexpression experiments. yFS1075 cells were first synchronized with nocodazole, then released into α-factor for a G1/S arrest, prior to release into S phase. Flask symbol indicates filtering and release into new media, with or without galactose and additives as indicated. **b)** Western blot of yFS1075 cells showing the levels of MCM at the indicated time points of the experiment, probed using anti-Mcm2-7 polyclonal antibody UM174. Noc = nocodazole arrest, α = α factor arrest. Boxed values indicate quantitation of Mcm2-7 levels in α-factor arrested cells with or without galactose induction, relative to Mcm2-7 levels of uninduced cells. **c)** Quantitation of flow cytometry data of the replicating population of yFS1075 cells as they progress from α-factor arrest through S phase. **d)** Normalized MNase-ChIP-seq coverage on Chromosome VIII for non galactose-induced cells at the α factor arrest point. ARS-annotated origins of replication are shown as black lines whereas unconfirmed origins are shown as orange lines/boxes.

### Increased MCM levels do not alter relative levels of helicase loading at origins

In order to measure MCM loading in cells overexpressing the full hexamer, we performed MNase ChIP-seq on α-factor arrested cells as outlined in **Figure 5a**. Fragments of the genome that were associated with MCM were pulled down, sequenced, and their abundance was normalized to an *S. pombe* spike-in control, resulting in peaks that correspond to previously annotated origins of replication (**Figure 5d**). In addition, the signal of MCM peaks correlates well with data from the MCM-depletion experiments, as well as previous publications (r = 0.78-0.93, **Figures S8a and S8b**).

To gauge whether increased cellular pools of MCM helicase lead to changes in loading during G1, we compared MCM abundance at origins of replication in cells overexpressing MCM and those that did not (**Table S1**). As seen in **Figure 6a**, MCM signal at origins in cultures overexpressing MCM show a ∼41% decrease in abundance compared to cultures with endogenous MCM levels. However, this difference is almost perfectly uniform throughout all origins assayed (r = 0.99), pointing to technical reasons involving ChIP or non-MCM-specific biological consequences of the overexpression. This result was not due to an insufficient amount of time allowed for loading, as three hours of MCM overexpression in α-factor produced similar loading levels (r = 0.99, **Figure S9a**). As a result, we conclude that overexpression of Mcm2-7 does not alter relative helicase loading in G1.

**Figure 6:**
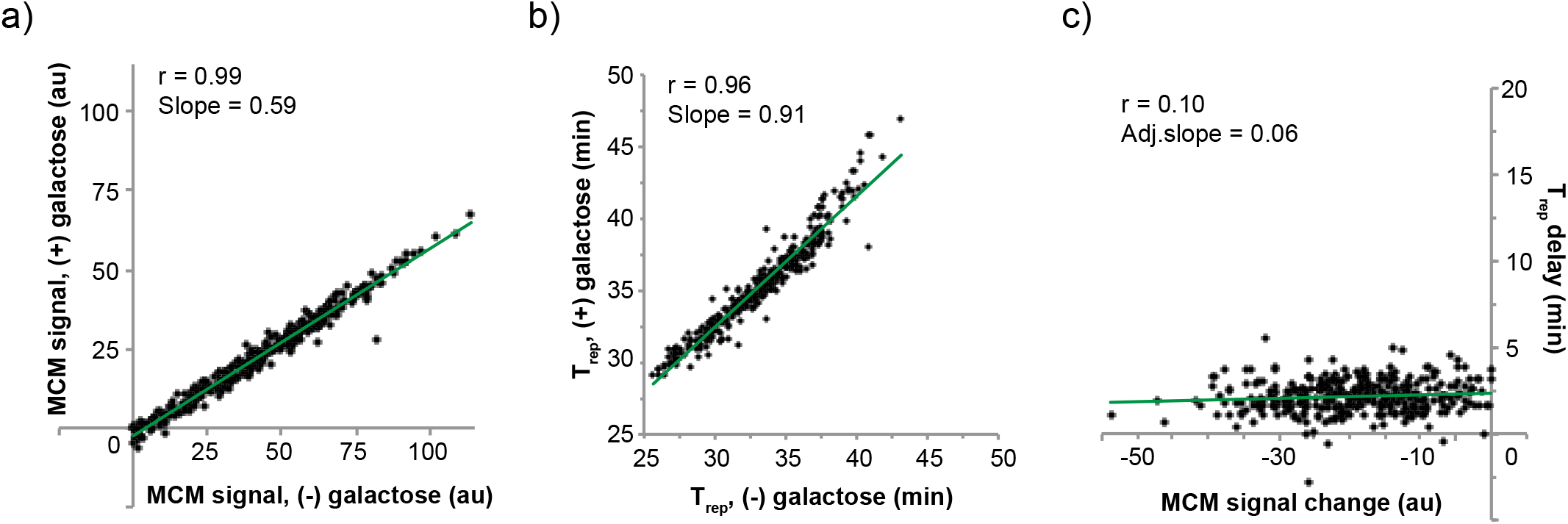
MCM overexpression does not cause changes to helicase loading or replication timing. **a)** yFS1075 cultures were synchronized and released into S phase as shown in Figure 5a. MNase-ChIP-seq signal at ARS-annotated origins at the α factor arrest point, with or without galactose-induced MCM overexpression. **b)** T_rep_ values for cultures released into S phase as in Figure 5a, with or without prior galactose-induced MCM overexpression. **c)** Correlation between delays in replication timing and changes in MCM signal as a result of galactose-induced MCM overexpression.

MCM requires Cdt1 to enter the nucleus and to be loaded onto origins of replication (Tanaka and Diffley, 2002). Although Cdt1 disassociates from the MCM complex after it is loaded onto DNA, it doesn’t fully exit the nucleus until late G1/early S phase. Therefore, although Cdt1 likely shuttles in and out of the nucleus in some equilibrium during the α-factor arrest in our experiments, it may not be able to accommodate nuclear import for all of the overexpressed MCM2-7 hexamers. To address the possibility that overexpressed MCM is not imported into the nucleus and therefore not loaded onto DNA due to a substoichiometric levels of Cdt1, we overexpressed MCM2-7 along with Cdt1. However, we did not observe any significant changes in MCM loading (r = 0.98, **Figure S9c**). Furthermore, the lack of any changes was not due to the inability of the overexpressed hexamer to enter cell nuclei, as microscopy confirmed the presence of overexpressed Mcm7 in cell nuclei (**Figure S10**). All together, these results indicate that overexpression of MCM helicase does not cause altered loading dynamics in G1.

### Overexpression of MCM does not affect replication timing

To test whether the increase in cellular MCM abundance led to changes in the replication timing program, we performed sync-seq experiments. T_rep_ values for cells expressing endogenous levels of MCM correlate well with values obtained from the MCM-depletion experiments as well as previously published data (r = 0.92, **Figure S8c**) (Müller et al., 2014). Moreover, origin replication times for cells overexpressing MCM do not significantly differ from control cells, beyond a brief and global delay attributed to the switch in carbon source (**Figures 6b, 5c**). Moreover, changes in replication timing do not correlate with changes in MCM signal (**Figure 6c**). Altogether, this data shows that overexpression of the MCM helicase does not affect replication timing.

## Discussion

A unifying model of the mechanisms involved in establishing replication timing in eukaryotes remains elusive. Numerous *cis-* and *trans*-acting factors have been shown to affect timing, but the mechanism by which they do is unclear (Rhind and Gilbert, 2013). Since replication timing is primarily determined by the timing of origin firing, these factors are thought to affect the timing of MCM activation. However, it has also been noted that regulating MCM occupancy at origins would affect replication timing, because higher MCM occupancy at an origin would lead to a higher probability of it firing and thus an earlier average firing time (de Moura et al., 2010; Yang et al., 2010). Variable MCM occupancy could occur in a scenario in which an origin can either be loaded with an MCM double-hexamer complex or not. In this single-MCM scenario, MCM occupancy at a specific origin is the fraction, from 0 to 1, of cells in which an MCM complex is loaded at that origin, a parameter that has been referred to as origin competency (de Moura et al., 2010). Alternatively, in a multiple-MCM scenario, more than one MCM double-hexamer complex can be loaded at a single origin, so MCM occupancy can range from 0 to many. Both single- and multiple-MCM scenarios have been proposed (Das et al., 2015; Foss et al., 2020; de Moura et al., 2010; Das and Rhind, 2016; Yang et al., 2010).

Our previous work supports a model in which multiple MCMs are loaded at origins (Das et al., 2015; Yang et al., 2010). Furthermore, overexpression of origin-activating factors in S phase causes most all origins to fire early in S phase, consistent with most origins having at least one MCM loaded (Lynch et al., 2019). Our MNase approach to MCM ChIP-seq reported here further supports the multiple-MCM model by producing high-resolution maps of MCM locations that identify MCMs at multiple locations across origins. We find MCM ChIP-seq signal distributed around origins up to three nucleosomes away from the ACS, similar to previous lower resolution results (Belsky et al., 2015), in addition to signal inside the origin NFR (**Figure 3a**). The location of MCMs in the NFR and the surrounding nucleosome support for a model in which MCMs are mobile after being loaded and can slide past nucleosomes during nucleosome exchange. Indeed, this data is consistent with observations that early origins, which tend to have more MCM signal by ChIP-seq (Das et al., 2015 and **Figure 4c**), display higher rates of nucleosome exchange (Dion et al., 2007). Furthermore, the fact that we find MCM with equal frequency upstream and downstream of origins suggests that ORC does not prevent MCM from diffusing past the ACS, consistent with ORC having a short dwell time at origins (Ticau et al., 2015; Sonneville et al., 2012). Nonetheless, because our current ChIP-seq data only represents an average of MCM locations, it cannot formally exclude the possibly that no more that one MCM double-hexamer complex is ever loaded at a single origin—albeit at varying locations—and thus our current data does not discriminate between the single- and multiple-MCM scenarios. However, for all the analyses presented here, the two scenarios are formally equivalent and make the same predictions about how competition between origins for MCM loading affects replication timing.

Regardless of stoichiometry, the possibility of variable MCM occupancy raises interesting questions about how MCM occupancy is regulated and how it affects replication timing. We considered two hypotheses for how varying MCM occupancy between origins could be regulated. The first hypothesis, which we refer to as the ORC Activity model, posits that the rate at which ORC loads MCM varies among origins. ORC activity at an origin could vary due to ORC occupancy, which could be affected by ORC affinity for the ACS or by the local chromatin environment (Hoggard et al., 2013), or due to ORC specific activity, which could be affected by local activators or repressors. The second hypothesis, which we refer to as the Origin Capacity model, posits that origins have an intrinsic capacity for MCM loading that varies among origins, and that once that capacity is achieved, no more MCMs can be loaded. To distinguish between these models, we varied cellular MCM levels, reasoning that changes in cellular MCM levels would lead to predictable changes in the level of MCM loaded at origins, depending on the mechanism that regulates origin loading. In particular, in response to a reduction in cellular MCM levels, the Origin Activity model predicts that all origins with be proportionally reduced in MCM occupancy, because origins will load at the same relative rates but the MCM pool will be depleted sooner, whereas the Origin Capacity model predicts that origins with low capacities will fill up as normal, but that origins with higher capacities will fail to reach those capacities once the MCM pool is depleted (**Figure 7a**).

**Figure 7:**
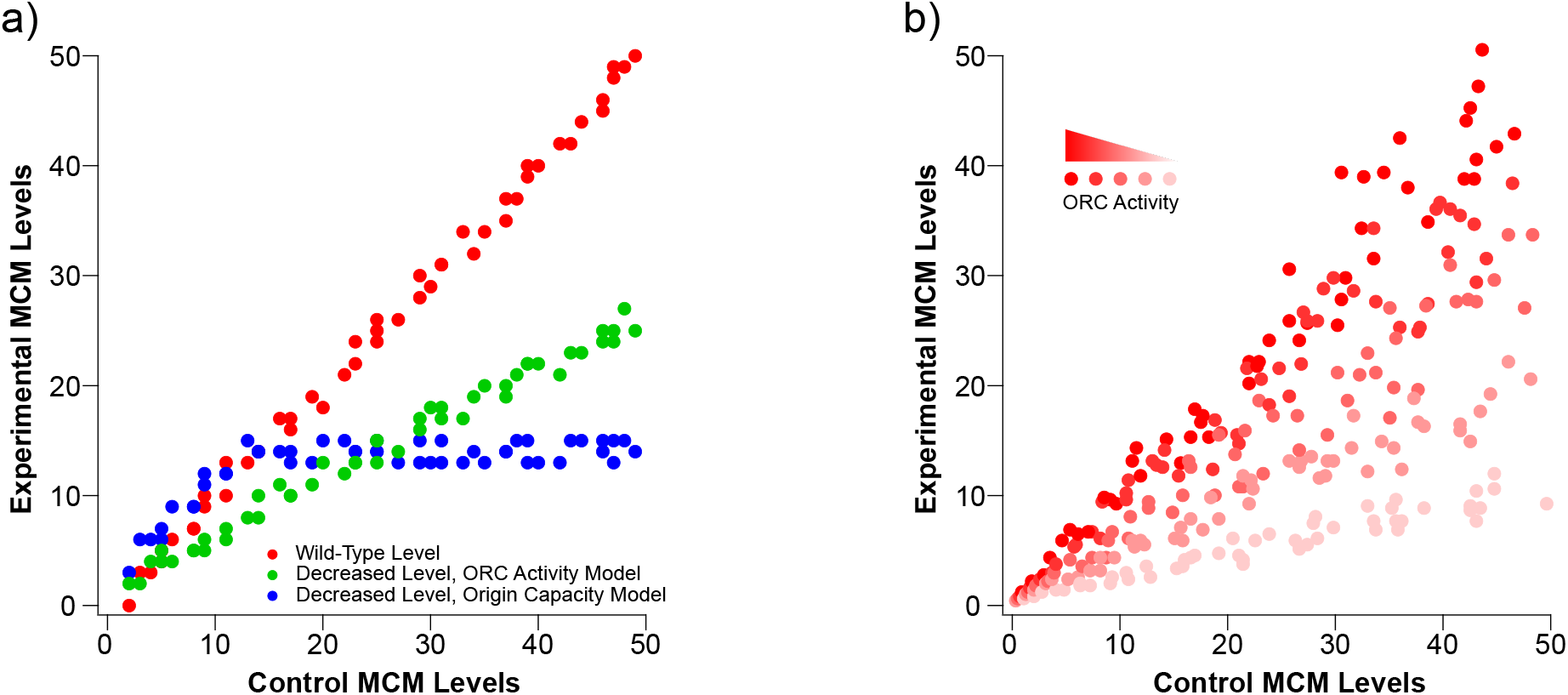
Models for how changes in the cellular levels of MCM affect their loading dynamics at origins of replication and replication timing. **a)** Expected results if an ORC Activity model or an Origin Capacity model governs how MCM is distributed genome-wide. **b)** A combined model shows that, if there is a range of ORC activities and origin capacities, in normal cells ORC activity has lile effect and MCM loading is largely determined by origin capacity because neither time nor MCM abundance are limiting. However, when MCM abundance becomes limiting, ORC activity becomes important because origins with more active ORC can load MCM faster.

Our results are not consistent with either the simple ORC Activity or Origin Capacity model (**Figures 2a and 7a**). Instead, we propose a hybrid model, in which both ORC activity and origin capacity vary at each origin (**Figure 7b**). In the hybrid model, origin capacity is the primary effector of MCM occupancy in wild-type cells. ORC activity does vary among origins, but because neither time during G1 nor MCM amount is limiting, all origins are loaded to their full capacity. However, when MCM levels become limiting, not all origins can be fully occupied. In this case, origins with high ORC activity (bright red in **Figure 7b**) outcompete other origins and are still loaded to their full capacity, whereas origins with progressively lower ORC activity (light red and pink in **Figure 7b**) load MCM to progressively lower proportions of their full capacity before the MCM pool is exhausted.

Our hybrid model is consistent with the results of our MCM-reduction experiments (**Figure 2a**) and, in fact, was motivated by the behavior of three classes of origins previously characterized by ORC affinity and origin activity (Hoggard et al., 2013 and **Figure 2c**). These three classes—DNA-dependent, chromatin-dependent and weak—show the behavior predicted by the hybrid model, with the strong, DNA-dependent origins maintaining close to full MCM loading (109-94%), the somewhat less strong, chromatin-dependent origins showing an intermediate reduction (100-77%), and the weak origins showing the greatest reduction in MCM loading (74-48%). Our hybrid model is also consistent with our MCM over-expression experiments. We find no change in the relative MCM ChIP-seq signal at origins in response to a five-fold increase in cellular MCM levels (r = 0.99, **Figure 6a**). This result is consistent with the conclusion that origins in wild-type cells are loaded to their full capacity and that MCM is normally in excess in budding yeast (Donovan et al., 1997).

Although we find no relative difference in MCM ChIP-seq signal in our MCM overexpression experiments, we find an absolute reduction of 59% (**Figure 6a**). This result was dependent on overexpression constructs being present in the cells, because a control strain with no overexpressing genes did not display any reduction in MCM ChIP-seq signal in response to galactose (**Figure S9b**). This result does not appear to be due to a uniform increase in background signal, which migth be expected if excess MCM was loaded with low specificity at many sites across the genome, because the data was normalized to an *S. pombe* spike-in to allow for absolute quantitation. It could be that the overexpression of MCM titrates away an interacting component that is crucial for MCM loading or nuclear import. However, in that case we would expect to see an effect similar to reducing MCM levels, which does not affect all origins uniformly (**Figure 2a**). Alternatively, although the antibody is added in saturating amounts, antibody-binding dynamics of chromatin-bound MCM could be altered due to the presence of excess MCM. In any case, given the lack of a plausible mechanistic explanation for reduced MCM loading in response to MCM overexpression, and the nearly perfect correlation between plus and minus overexpression conditions, we conclude that increased MCM levels do not cause altered MCM loading in yeast.

In addition to the lack of changes in MCM loading, overexpression of MCM did not affect cell viability, bulk S-phase progression, or replication timing (**Figures 6**). In contrast, overexpression of MCM along with Cdt1, but not Cdt1 alone, was lethal if constitutively expressed (**Figure S6a**), resulting in dumbbell-shaped terminal phenotype and suggesting a metaphase arrest. This result does not appear to be related to MCM loading or replication timing (at least in the window of one cell cycle) as levels of MCM loading and replication timing remained largely unchanged in MCM/Cdt1-overexpressing cells (**Figure S8c**). Although this points to a synergistic effect between MCM and Cdt1 in causing cell death, the exact mechanism for the lethality remains unclear.

Reduction of MCM levels caused a delay in progression through bulk S phase (**Figures 1c and 4a**). The slower progression through S phase is consistent with less MCM being loaded onto some origins, resulting in fewer origins being activated in S phase. If MCM levels at all origins were uniformly reduced, we would not expect any change in replication timing because replication initiation in budding yeast is regulated by competition for a limited a set of initiation factors (Mantiero et al., 2011; Tanaka et al., 2011). If MCM loading were lowered in uniform proportions, the relative competition for initiation factors, and thus replication timing, would be maintained. However, because the reduction of MCM loading is not uniform, and because low MCM origins tend to lose more MCM than high efficiency ones (**Figure 2b**), the distribution of MCM is more heterogeneous, leading to larger distances between efficient origins and thus longer S phase (**Figure S3, Table S2**).

Our combination of MCM ChIP-seq and replication timing data also allow us to directly confirm in a single experiment previous conclusions, drawn from comparing data collected in different studies, that replication timing correlates with levels of MCM loading (Das et al., 2015; Foss et al., 2020). Although the correlation between MCM levels and replication timing is robust (**Figure 4c**, p > 10^−5^), it is clearly not the only determining factor (r = 0.29-0.52). Therefore, we conclude that MCM loading is a significant effector of replication timing, but one that is modulated by other factors that affect the specific activity of loaded MCMs. Nonetheless, by manipulating MCM levels and showing that replication changes concomitantly, we confirm genome-wide that MCM levels directly regulate replication timing (**Figure 4d**), an experiment that had previously been done only at a single locus (Das et al., 2015).

In this study, we show that some origins are efficient at recruiting MCM, while others depend on excess levels of the helicase to display the loading observed in wild type cells. These results indicate that the relative levels of MCM loaded at origins are not simply static, but rather a dynamic balance dependent on cellular MCM levels, origin MCM recruiting ability and origin MCM capacity. In addition, our data suggests that wild-type MCM abundance is saturating, such that increasing levels further does not lead to increased loading. Altogether, this work suggests that MCM loading homeostasis, and subsequently replication timing, in wild type cells is dependent on MCM being in excess. In future studies, it will be interesting to dissect the specific *cis*- and *trans*-acting factors that make origins sensitive or resistant to changes in MCM levels.

## Materials and Methods

Strains used are listed in Table 1. Standard techniques were used for strain constructions. Cultures were grown in YP-Dextrose (YPD) at 30°C, unless otherwise noted.

**Table 1.** Strains used

| Strain | Genotype | Source |
| --- | --- | --- |
| yFS105 | h- leu1-32 ura4-D18 | Lab WT *pombe* strain |
| yFS833 | $\alpha$ leu2-3,112 ura3 trp1-1 ade2-1 can1-100 his3-11,15 GAL+ ssd1-d2 | (Müller et al., 2014) |
| yFS1020 | a leu2-3,112 ura3-1 trp1-1 ade2-1 can1-100 his3-11,15 bar1-Δ lys2::HisG Pep4Δ | (Ticau et al., 2015) |
| yFS1021 | a leu2-3,112 ura3-1 trp1-1 ade2-1 can1-100 his3-11,15 bar1::HisG lys2::HisG Pep4Δ Trp1::Gal1,10-MCM6,MCM7 His3::Gal1,10-MCM2,MCM3 Lys2::Gal1,10-MCM4,MCM5 Ura3::Gal1,10-Cdt1,Gal4 | (Ticau et al., 2015) |
| yFS1044 | a leu2-3,112 ura3-1::ADH1-osTIR1-2-9myc (URA3) trp1-1 ade2-1 can1-100 his3-11,15 | (Nishimura et al., 2009) |
| yFS1059 | a leu2-3,112 ura3-1::ADH1-osTIR1-2-9myc (URA3) trp1-1 ade2-1 can1-100 his3-11,15 MCM4::MCM4-IAA17-GFP(HPH) | This study |
| yFS1062 | a his3-11,15 leu2-3,112 ura3-1::ADH1-osTIR1-2-9Myc(URA3) trp1-1 ade2-1 can1-100 Mcm4::Mcm4-IAA17(kanMX) | This study |
| yFS1075 | a leu2-3,112 ura3-1 trp1-1 ade2-1 can1-100 his3-11,15 bar1-Δ lys2::HisG Pep4Δ Trp1::Gal1,10-MCM6,MCM7-GFP His3::Gal1,10-MCM2,MCM3 Lys2::Gal1,10-MCM4,MCM5 | This study |
| yFS1076 | a leu2-3,112 ura3-1 trp1-1 ade2-1 can1-100 his3-11,15 bar1-Δ lys2::HisG Pep4Δ Trp1::Gal1,10-MCM6,MCM7 His3::Gal1,10-MCM2,MCM3 Lys2::Gal1,10-MCM4,MCM5 | This study |
| yFS1080 | a leu2-3,112 ura3-1 trp1-1 ade2-1 can1-100 his3-11,15 bar1-Δ lys2::HisG Pep4Δ Ura3::Gal1,10-Cdt1,Gal4 | This study |
| yFS1081 | a leu2-3,112 ura3-1 trp1-1 ade2-1 can1-100 his3-11,15 bar1-Δ lys2::HisG Pep4Δ Ura3::Gal1,10-Cdt1,Gal4 Trp1::Gal1,10-MCM6,MCM7 His3::Gal1,10-MCM2,MCM3 Lys2::Gal1,10-MCM4-GFP,MCM5 Ura3::Gal1,10-Cdt1,Gal4 | This study |
| yFS1082 | a leu2Δ0 ura3Δ0 his3Δ1 met15Δ0 MCM4-GFP(his3mx6) | (Huh et al., 2003) |
| yFS1083 | a leu2Δ0 ura3Δ0 his3Δ1 met15Δ0 MCM7-GFP(his3mx6) | (Huh et al., 2003) |

### Auxin degradation experiments

*S. cerevisiae* strain yFS1059 (MCM4-AID-GFP) was grown in YPD at 30°C to an OD of ∼0.2. Nocodazole was added to a final concentration of 7.5 µg/ml for two hours. 1.5 hours into the nocodazole arrest, cultures were supplemented with auxin (Indole-3-acetic acid sodium salt, Sigma I5948) to final concentrations of 500 µM, 30 µM, or 0 µM. After two hours total in nocodazole, cultures were vacuum-filtered and cells were resuspended in fresh YPD supplemented with 5 µg/ml α-factor (Biomatik) and auxin to a final concentration of 500 µM, 30 µM, or 0 µM, as before filtration. After 1.5 hours, cultures were vacuum filtered and cells were resuspended in fresh YPD supplemented with 500 µM, 30 µM, or 0 µM, as before filtration, for 30 minutes. Cultures were vacuum-filtered a final time and cells were resuspended in fresh YPD for S phase progression.

For ChIP sample collection, α-factor-arrested cells were crosslinked with formaldehyde and collected as previously described (Wal & Pugh 2012).

### Galactose overexpression experiments

*S. cerevisiae* strains yFS1075 (GAL-MCM), yFS1021 (GAL-MCM, GAL-CDT1), and yFS1020 (WT) were grown in YP-Raffinose (YP-Raf) at 30°C to an OD of ∼0.2. Nocodazole was added to a final concentration of 10 µg/ml. After two hours, cultures were vacuum-filtered and cells were resuspended in fresh YP-Raf supplemented with α-factor (Biomatik) to a final concentration of 25 nM, as well as either galactose to a final concentration of 2% or an equal volume of water. Cells were allowed to arrest in α-factor for either two or three hours, followed by release via vacuum filtration into fresh YP-Dextrose (YPD) supplemented with 0.2 mg/ml pronase (Sigma P6911).

For ChIP sample collection, α-factor arrested cells were crosslinked with formaldehyde and collected as previously described (Wal & Pugh 2012). After formaldehyde quenching, *S. cerevisiae* cells were combined with log-phase wild-type *S. pombe* cells (yFS105, also crosslinked) at a 9:1 ratio.

### Fluorescence microscopy

Cultures were grown and supplemented with 2% final galactose as in Figure 5a. At the α-factor-arrest point, 1ml cells were fixed by the addition of 10ml methanol which was pre-chilled in dry ice, then stored at −80°C until imaging. To prepare for imaging, fixed cells were equilibrated to 4°C from their −80°C storage, centrifuged, and all but 1ml supernatant was removed. The remaining mix was washed 2x with 2 ml ice cold PBS. Cells were then resuspended in 150 µl Vectashield+DAPI (Vector Laboratories H-1200) and imaged using a Zeiss Axioskop 2 Plus microscope fitted with DIC, GFP, and DAPI filters.

### Spot Assays

Yeast was grown in YPD (auxin strains) or YP-Raf (Galactose-overexpression strains) overnight at 30°C. Log phase cells were collected by brief centrifugation, washed twice with PBS, serially diluted five-fold starting with 4×10^6^ cells, then spotted onto the specified medium. Plates were incubated at 30°C for at least 36 hours before imaging.

### Western Blots and Quantitations

Standard western blot techniques were used (Green, 2012). Mcm2-7 was probed with polyclonal UM174 (gift from Bell lab, MIT - (Chen and Bell, 2011)) and anti-rabbit HRP conjugate secondary (Promega W401B), tubulin was probed with monoclonal anti-tubulin antibody (Sigma T5168) and anti-mouse HRP conjugate secondary (Promega W402B), and GFP was probed with the monoclonal JL8 antibody (Takara) and anti-mouse HRP conjugate secondary (Promega W401B). Probing of Mcm2-7 using UM174 antibody was performed using 12% gels in order to compress signal from the six MCMs into a compact region to facilitate quantification. Signal was detected using SuperSignal West Dura chemiluminescent HRP substrate (Thermo 34076) and GE Amersham Imager 600. Quantitation of gel images was carried out using ImageJ (NIH, Maryland, USA).

### ChIP Experiments

ChIP experiments were carried out as described previously with modifications noted below (Wal & Pugh 2012). MNase was titrated for each sample to determine concentrations that would yield ∼80-90% mononucleosomal digestion. Immunoprecipitation was carried out using 95% of the MNase digested sample (v/v) and 4 µl of anti-MCM2-7 polyclonal antibody (UM174, gift from Bell lab, MIT - (Chen and Bell, 2011)). The remaining 5% of MNase digested sample was processed as ‘input’ and was treated with Proteinase K and RNase A to prepare libraries for deep sequencing.

### Flow Cytometry

0.25 ODs of fixed cells (see below) were washed once with water, resuspended in 250 µl RNase A solution (100 µg/ml RNase A, 50 mM Tris pH 8.0, 15 mM NaCl), and incubated overnight at 37°C. Cells were then collected and resuspended in Proteinase K solution (125 µg/ml Proteinase K, 50 mM Tris pH 8.0) for 1 hour at 50°C. The samples were then collected by centrifugation, resuspended in 1 ml staining solution (1 µM Sytox green –ThermoFisher S7020, 50 mM Tris pH8.0), sonicated briefly using a microtip sonicator, and analyzed for flow cytometry using a Guava easyCyte instrument.

### Sync-seq Replication Timing Experiments

Experiments to monitor replication timing consisted of measuring population movement through S phase by flow cytometry and measuring genome-wide replication timing by deep sequencing (Müller et al., 2014). To do so, 3 ODs of cells were collected for each time point and arrested by the addition of sodium azide to 0.1% and EDTA to 20mM. From these samples, 0.25 ODs were used for flow cytometry and the rest was used for genome-wide copy number analysis by deep sequencing.

To prepare genomic DNA for deep sequencing, cells were lysed as for ChIP experiments. Lysates were treated with proteinase K and RNase A, followed by two consecutive extractions with phenol-chloroform-IAA (25:24:1, Fisher). DNA was purified by ethanol precipitation and resuspended in 135 µl water in preparation for shearing. A Covaris machine was used to shear the DNA to an average size of ∼200bp, following manufacturer’s instructions. DNA was then purified by DNA Clean and Concentrator Columns (Zymo Research) before making libraries.

### Library Preparation and Sequencing

Libraries were prepared using Next Ultra II kits (NEB) following the manufacturer’s protocol. Following library preparation, ChIP samples were purified as described (Wal and Pugh, 2012), while input samples were purified using a 2:1 ratio of AmpureXP (GE) beads to library. Following PCR amplification, all samples were purified using AMpure XP beads with 0.9:1 beads to PCR ratio. Samples were sequenced using the Illumina NextSeq 500 platform with paired ends.

### Data Analysis

The sequencing data from this study have been submitted to the NCBI Sequence Read Archive under accession number PRJNA663099. For MNase-ChIP experiments, sequencing reads were mapped to SacCer3 using Bowtie1, a 650bp upper limit cutoff, and exclusion of reads mapping to more than one genomic location. Duplicates were removed using Samtools 1.4.1. ACS-centered density profiles were constructed using coverage files normalized as indicated in the paragraph below. V plot intensities were normalized to counts per million (CPM) and generated by taking into account strand specificity of the ACSs (Eaton et al., 2010) using R and modified in-house scripts (courtesy of Nils Krietenstein). Heatmaps were generated using Deeptools 3.0.2 without taking strand specificity into account. Quantification of MCM signal at origins of replication was done using a 1 kb window centered at the origin or ACS midpoints. Correlation coefficients and slopes were calculated for origins displaying normalized MCM signal greater than zero for untreated cells. The set of origins displaying ARS activity in plasmid assays from OriDB (Siow et al., 2012) was used for all quantitations, unless otherwise stated. Replicate #1 was used for all analyses except for V plots, for which the higher resolution Replicate #2 was used.

Coverage files for auxin degradation experiments were normalized to CPM and to the non-origin background signal in 25 bp windows. Input signal was then subtracted from the normalized ChIP coverage files to account for MNase digest-induced background oscillations. For galactose induction experiments, ChIP coverage files were normalized in 25 bp windows to the total number of reads mapped to the *S. pombe* spike-in control. Normalized input signal was then subtracted from the normalized ChIP coverage to account for background oscillations in the data.

For replication timing experiments, coverage files were generated using LocalMapper scripts and T_rep_ was calculated in 1000 bp bins genome wide using Repliscope. Both packages were shared by Dzmitry Batrakou of the Nieduszynski Lab and are available at https://github.com/DzmitryGB/ (Müller et al. 2014). T_rep_ data generated from Repliscope was LOESS-smoothed in windows of ∼50kb using Igor (Wavemetrics, Lake Oswego, OR, USA). T_rep_ data for subtelomeric origins residing within 15 kb of chromosome ends was discarded due to large amount of noise. At least seven samples covering S phase progression were used to estimate T_rep_.

---

**Supplemental Figure 1:**
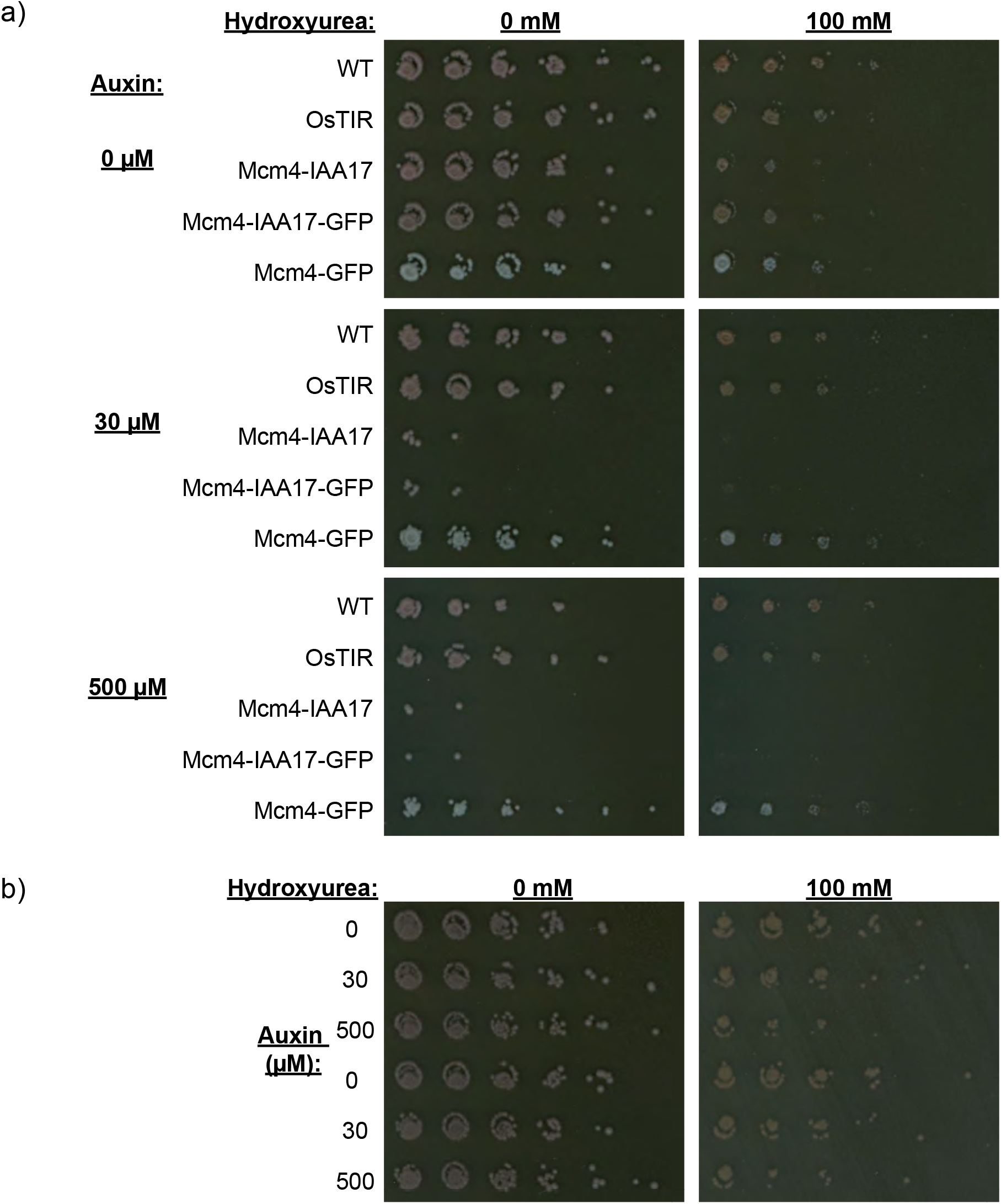
Auxin-induced degradation of Mcm4 causes reduced viability. **a)** For chronic down-regulation of MCM, exponentially-growing cultures of the indicated genotypes were serially diluted and spoed on plates containing 0 µM, 30 µM, or 500 µM auxin, with or without 100 mM hydroxyurea. OsTIR (yFS1044) refers to the background of the strain required for auxin-mediated degradation. WT = yFS833; MCM4-IAA17 = yFS1062; MCM4-IAA17-GFP = yFS1059; MCM4-GFP = yFS1082. **b)** For acute down-regulation of MCM, yFS1059 cultures were arrested at G1/S using α-factor as outlined in Figure 1a, washed, then serially diluted on YPD plates with or without 100 mM HU.

**Supplemental Figure 2:**
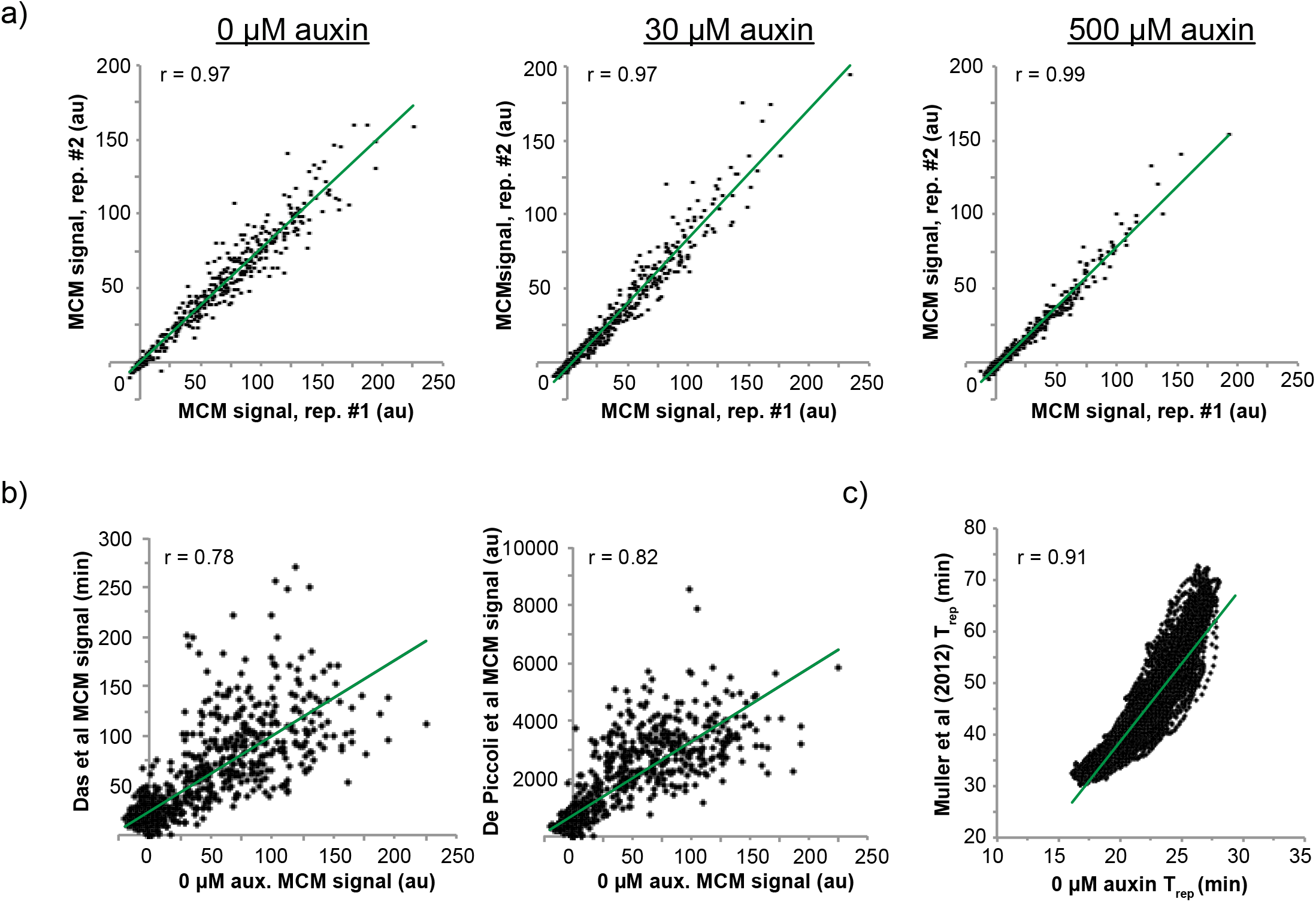
MNase-ChIP-seq and replication timing results from the MCM reduction experiments are reproducible. **a)** MCM ChIP-seq signal at ARS origins in two biological replicates for 0 µM, 30 µM, and 500 µM auxin treatments (yFS1059 strain). **b)** Comparison of MCM signal at origins between the 0 µM auxin condition in this study and other publications (Das et al., 2016, DePicoli et al., 2012). **c)** Genome-wide replication timing correlation in 1 kb windows between the 0 µM auxin condition in this study and Mueller et al, 2012.

**Supplemental Figure 3:**
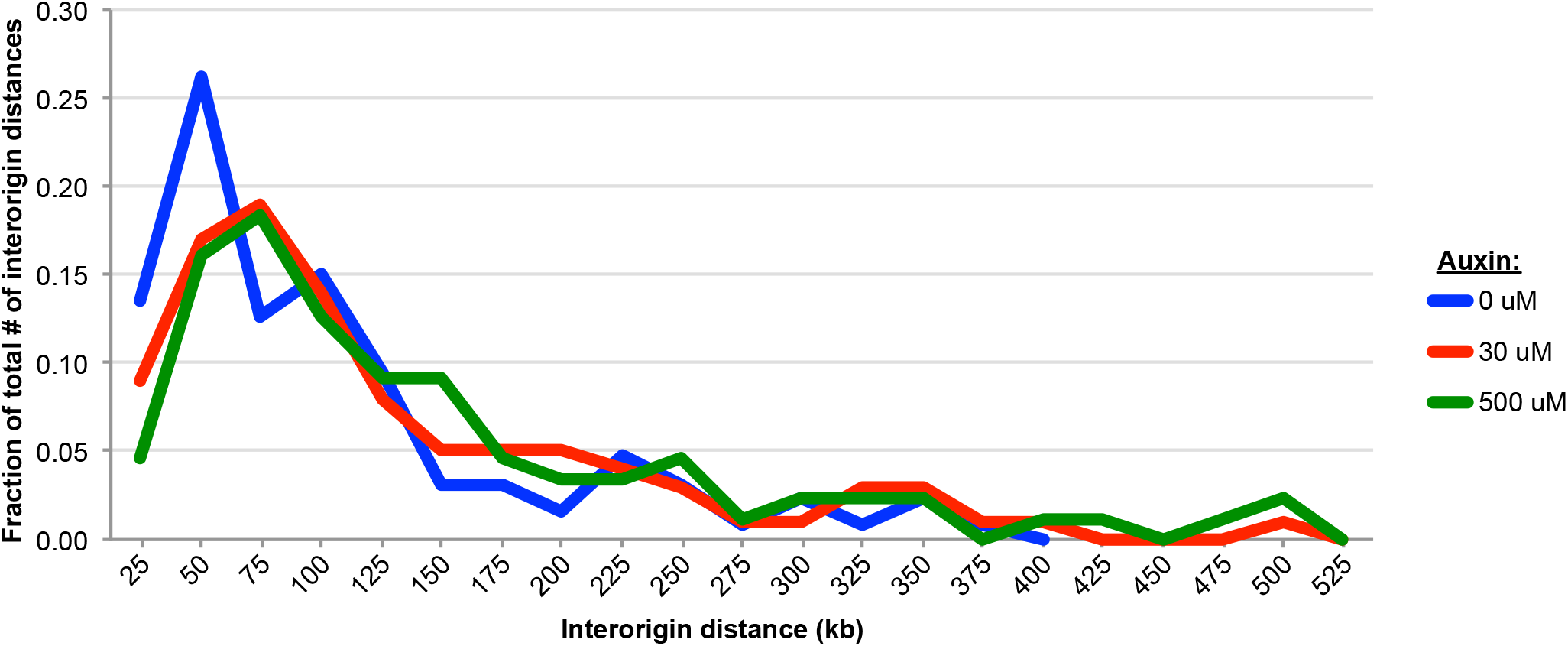
Genome-wide reduction in MCM loading leads to larger inter-origin distances. **a)** The smallest number of origins required to make up 50% of the total MCM signal was calculated for each condition. The inter-origin distances for these sets of origins were calculated and ploed as a fraction of the total number of inter-origin distances for each condition.

**Supplemental Figure 4:**
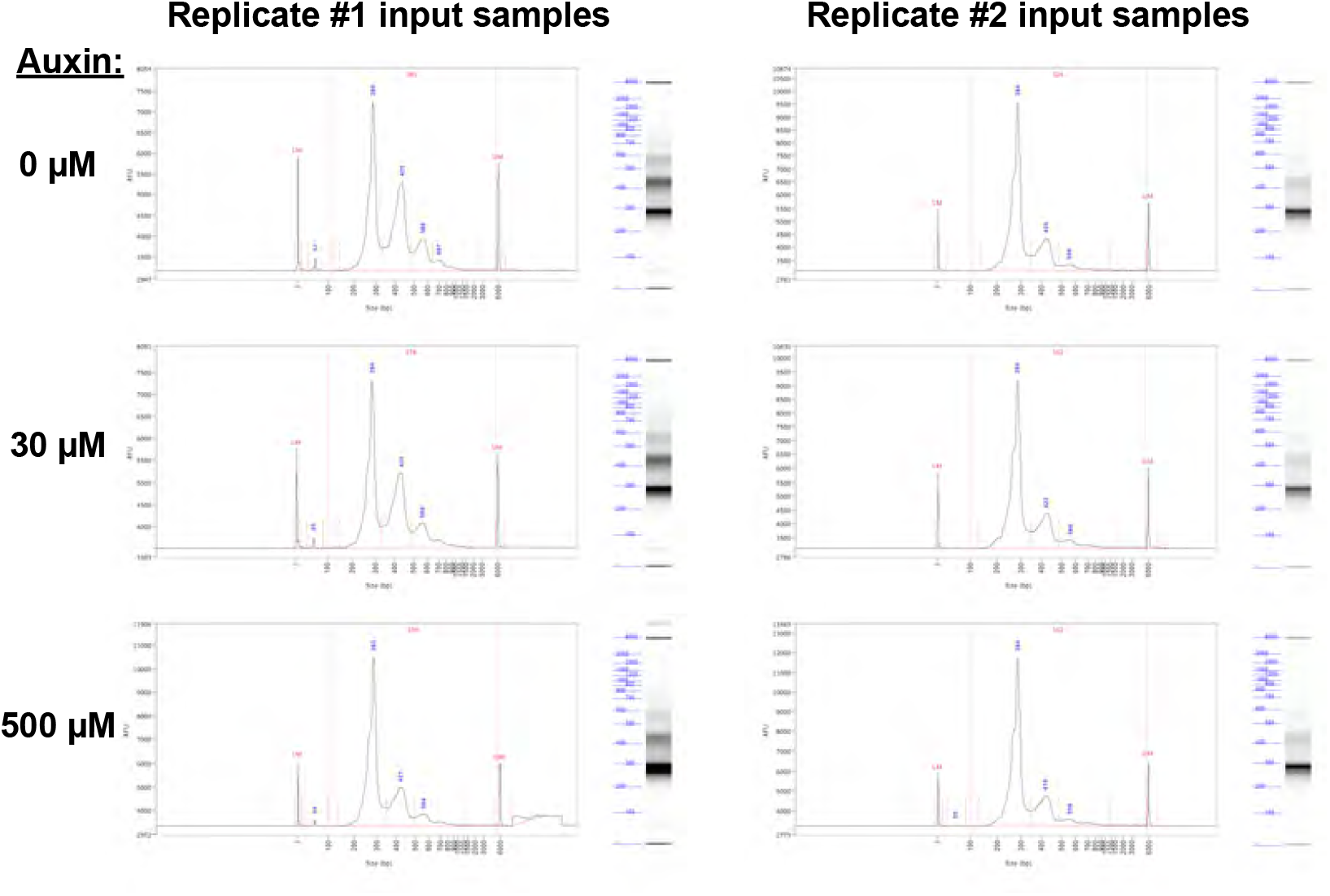
Fragment analyzer results for the two MCM reduction replicates shows different degrees of MNase digestion. Fragment Analyzer (Advanced Analytical Technologies) results of sequencing-ready input libraries show the different degrees of digestion for Replicates #1 and #2 (yFS1059 strain). Replicate #2 displays higher mononucleosome-sized (146bp nuc-fragment + sequencing adapters and primers = ∼284bp) content and therefore a stronger digestion.

**Supplemental Figure 5:**
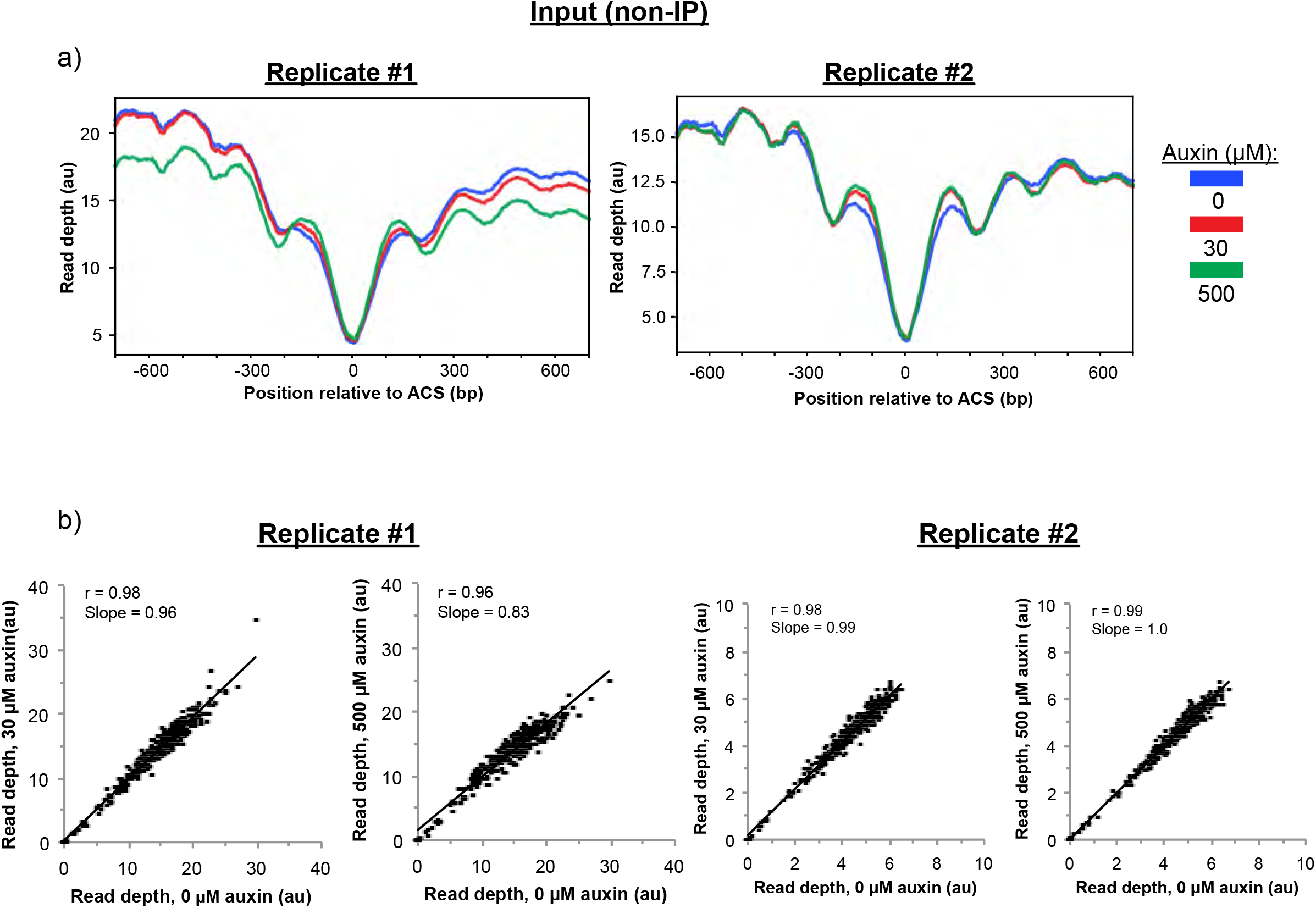
Input density profiles and origin coverage quantitations do not change significantly after auxin-induced Mcm4 degradation. **a)** Input (non-IP) read coverage density profiles at ARS origins of replication for Replicates #1 and #2 for the indicated auxin treatments (yFS1059 strain). **b)** Comparison of input coverage within 1 kb of ARS origins between 0 µM versus 30 µM and 500 µM auxin treatments.

**Supplemental Figure 6:**
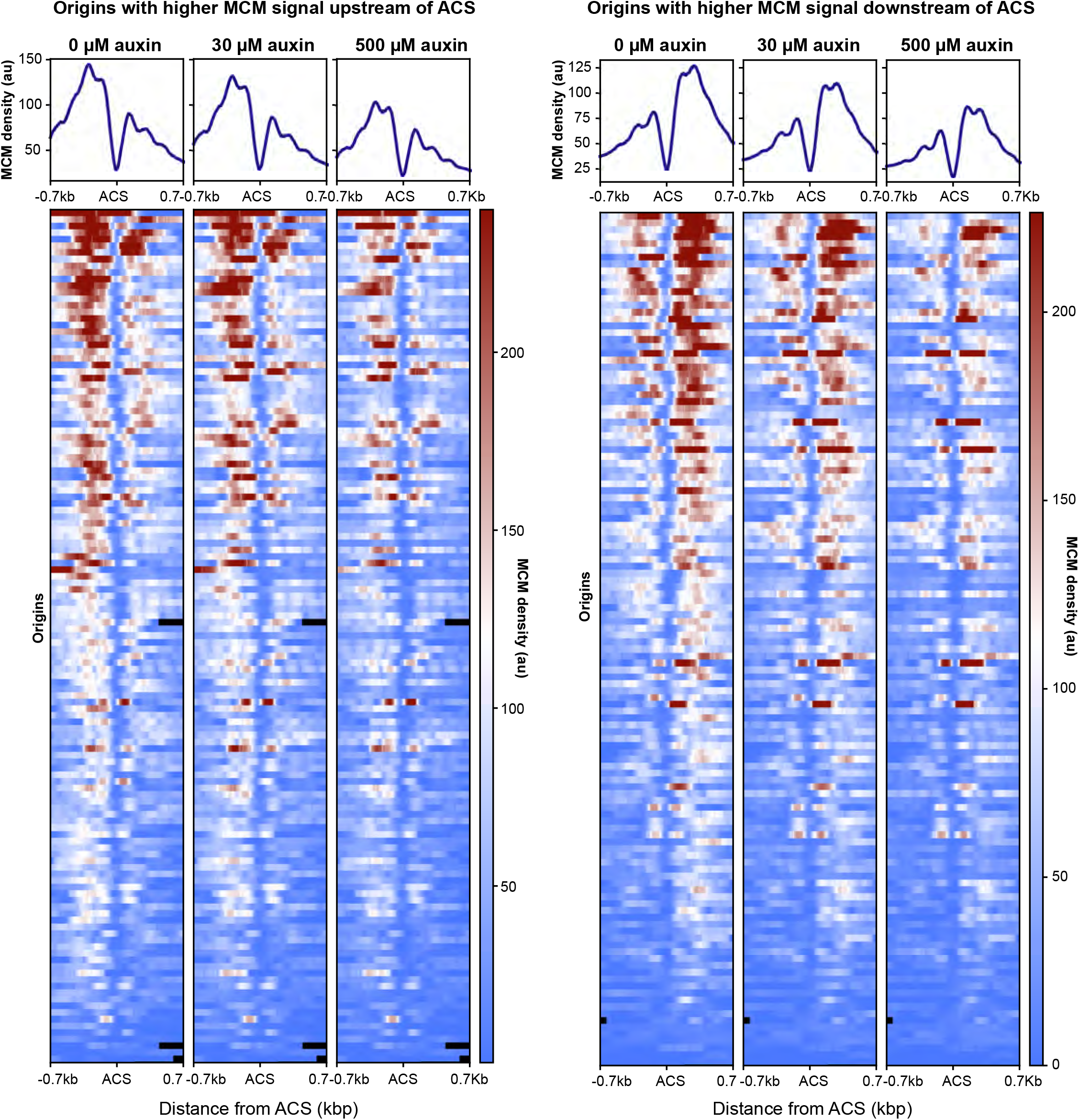
ACS-annotated origins separated by magnitude of signal upstream or downstream of ACS. MCM ChIP-seq signal was quantified 500 bp upstream and downstream of ACSs (yFS1059 strain). Origins were separated based on the dominant signal. Heatplots were generated for the set of origins displaying higher MCM signal upstream of the ACS as well as for the set of origins displaying higher signal downstream of the ACS.

**Supplemental Figure 7:**
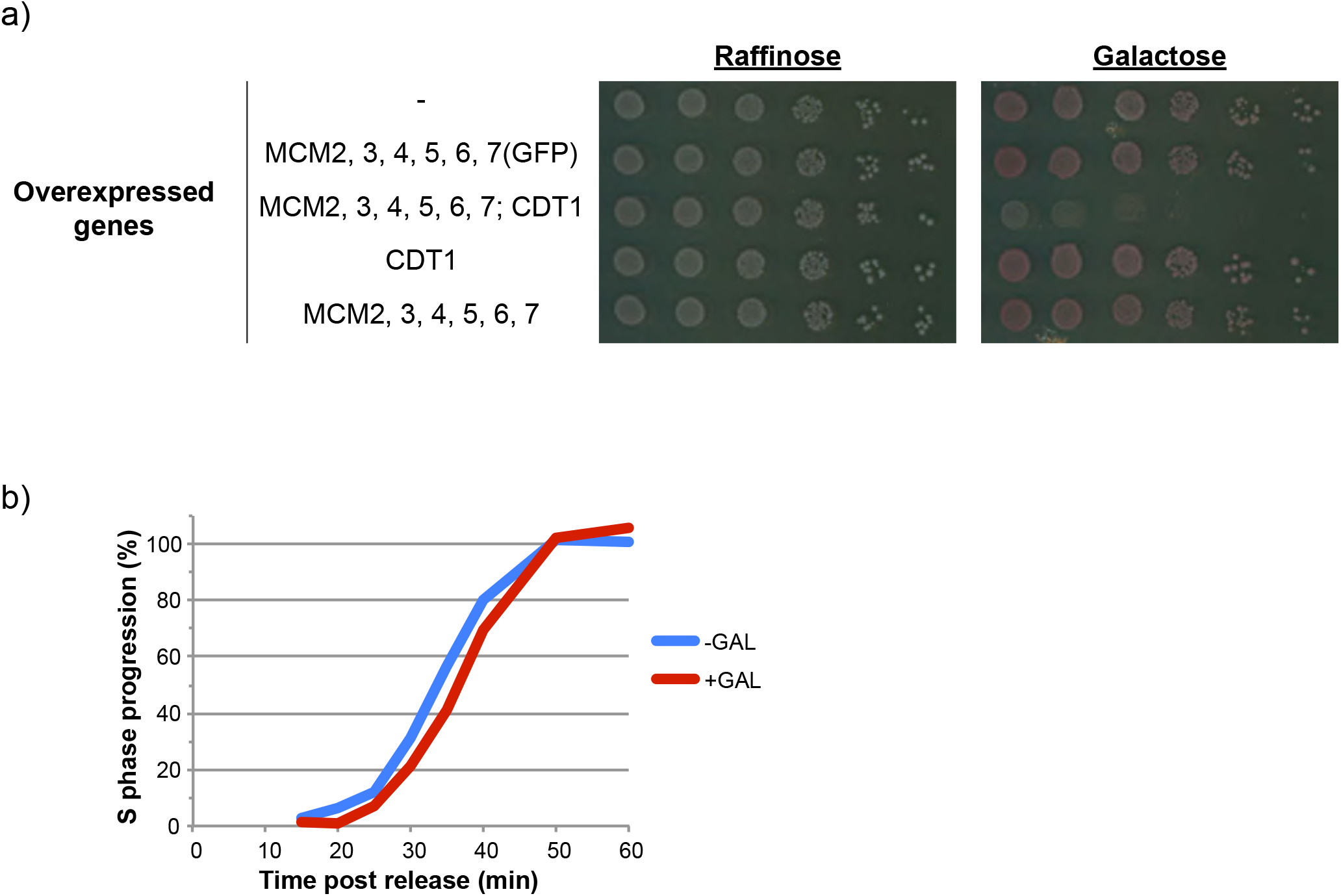
MCM overexpression does not affect cell viability. **a)** Serial dilution assay of exponentially growing cultures of the indicated overexpressing genotypes, grown in the presence of raffinose or galactose. (-) = yFS1020, MCM2,3,4,5,6,7(GFP) = yFS1075; MCM2,3,4,5,6,7, CDT1 = yFS1021; CDT1 = yFS1080; MCM2,3,4,5,6,7 = yFS1076. **b)** Cultures from a control strain (no overexpression vectors – yFS1020) were treated as in Figure 5a and released into S phase. Flow cytometry quantitation shows the S phase progression of cells supplemented with glucose during the α-factor arrest versus those there were not.

**Supplemental Figure 8:**
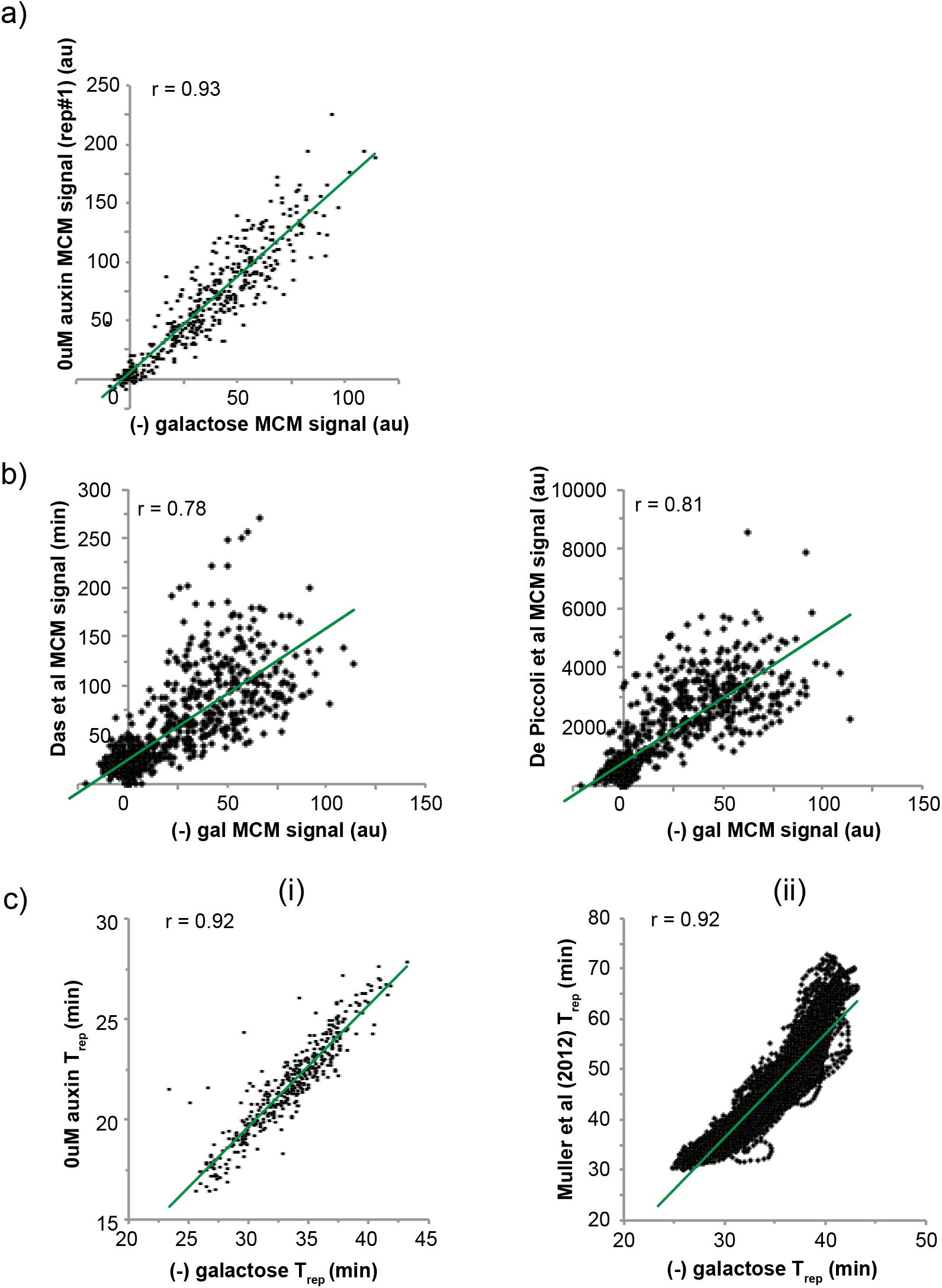
MNase-ChIP-seq and replication timing results from the MCM overexpression experiments are reproducible. **a)** MCM ChIP-seq signal at ARS origins for the two main ‘untreated’ samples in this study: 0 µM auxin condition (Figure1a – yFS1059) versus (-) galactose condition (Figure 5a – yFS1075). **b)** Comparison of MCM signal at origins between the (-) galactose condition in this study (yFS1075 strain) and other publications (Das et al., 2016, DePicoli et al., 2012). **c)** Replication timing correlation between the (-) galactose condition (yFS1075 strain) and: i) 0 µM auxin condition for ARS origins, and ii) genome-wide 1 kb windows from Mueller et al, 2012.

**Supplemental Figure 9:**
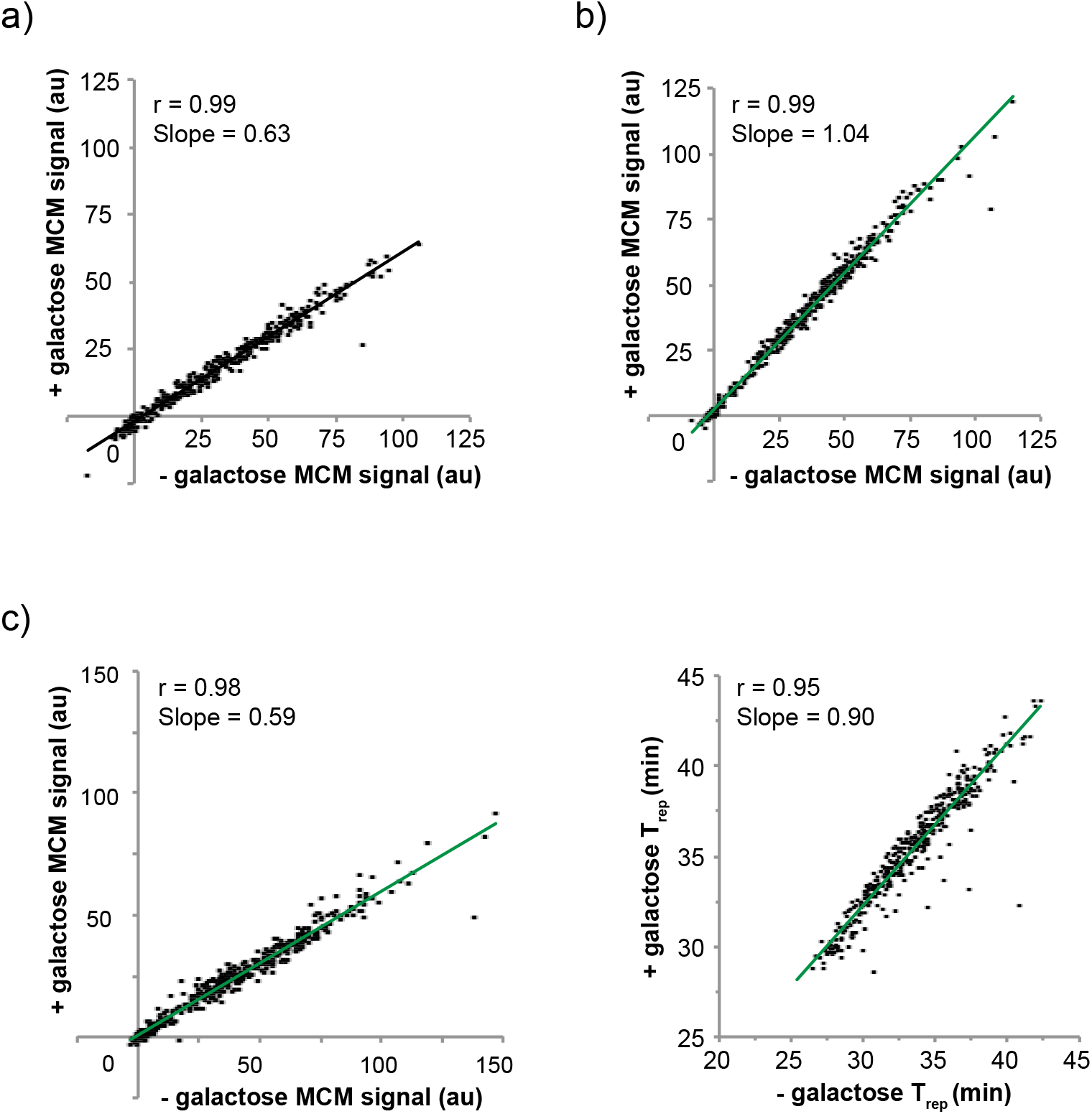
Overexpression of MCM for longer times or along with Cdt1 does not alter loading levels at origins or replication timing. **a)** Quantitation of MCM signal at ARS origins when MCM is overexpressed as in Figure 5a but for 3 hours (yFS1075). **b)** Quantitation of MCM signal at ARS origins for a control strain that does not overexpress any genes (yFS1020). **c)** Quantitation of MCM signal and T_rep_ at ARS origins when MCM is overexpressed along with Cdt1 as in Figure 5a (yFS1021).

**Supplemental Figure 10:**
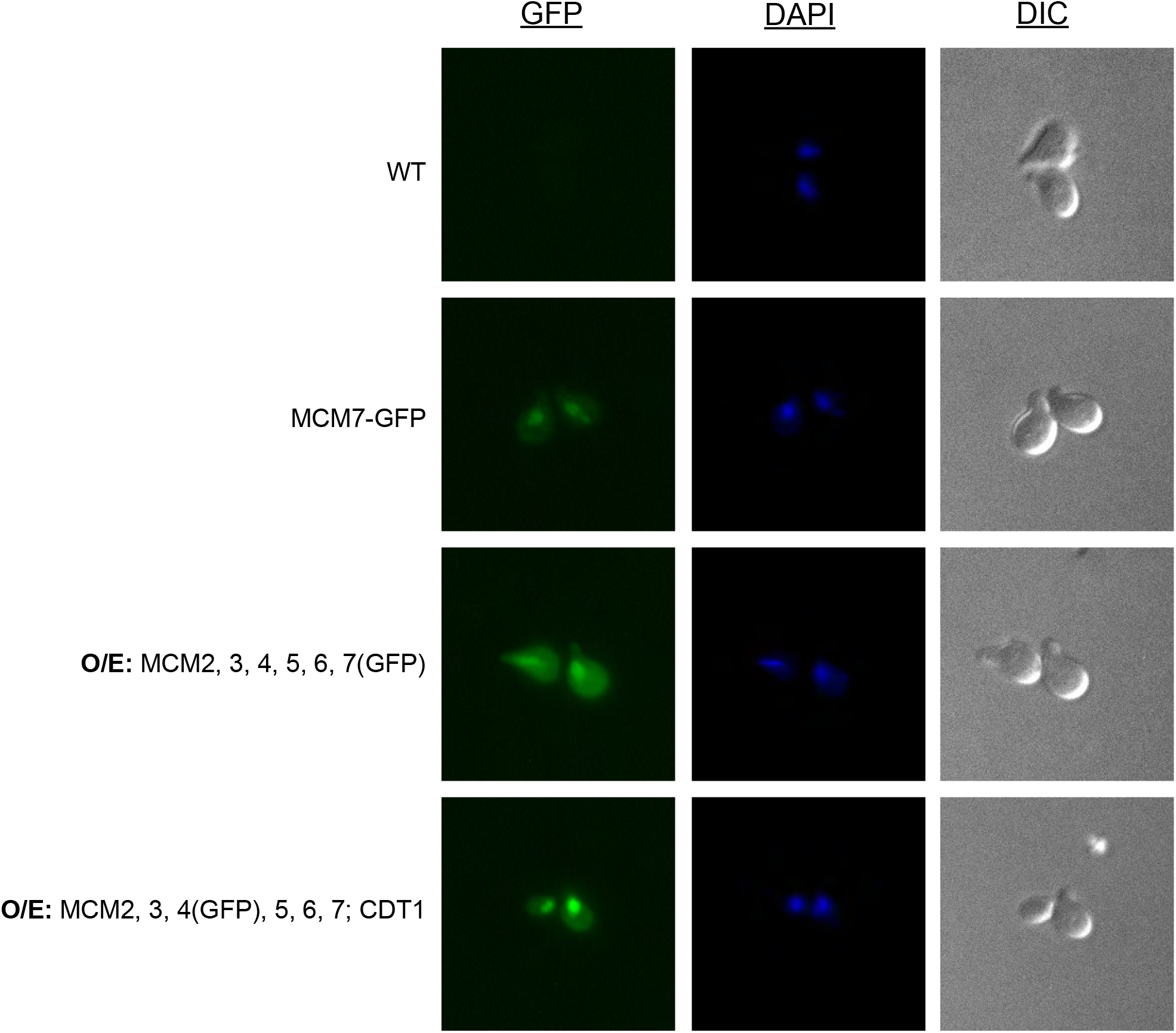
Overexpressed MCM is able to enter the nucleus during α-factor arrest. WT (yFS1020), MCM7-GFP (yFS1083), MCM2, 3, 4, 5, 6, 7-GFP overexpressing (yFS1075) and MCM2, 3, 4-GFP, 5, 6, 7 and CDT1 overexpressing (yFS10810 cultures were grown as in Figure 5a in the presence of galactose and visualized at the α factor arrest time point using DAPI to stain DNA and GFP fluorescence to visualize MCM quantity and localization.

## Notes

### Competing Interest Statement

The authors have declared no competing interest.

